# High Throughput pMHC-I Tetramer Library Production Using Chaperone Mediated Peptide Exchange

**DOI:** 10.1101/653477

**Authors:** Sarah A. Overall, Jugmohit S. Toor, Stephanie Hao, Mark Yarmarkovich, Son Nguyen, Alberto S. Japp, Danai Moschidi, Michael R. Betts, John M. Maris, Peter Smibert, Nikolaos G. Sgourakis

## Abstract

Peptide exchange technologies are essential for the generation of pMHC-multimer libraries, used to probe highly diverse, polyclonal TCR repertoires. Using the molecular chaperone TAPBPR, we present a robust method for the capture of stable, empty MHC-I molecules which can be readily tetramerized and loaded with peptides of choice in a high-throughput manner. Combined with tetramer barcoding using multi-modal cellular indexing technology (ECCITE-seq), our approach allows a combined analysis of TCR repertoires and other T-cell transcription profiles together with their cognate pMHC-I specificities in a single experiment.

## MAIN TEXT

T-cells recognize foreign or aberrant antigens presented by MHC-I expressing cells through the T-cell receptor (TCR) and is the first critical step towards establishment of protective immunity against viruses and tumors^1^. Fluorescently tagged, multivalent MHC class-I reagents (multimers) displaying individual peptides of interest have revolutionized detection of antigen specific T-cells^2^. Staining with multimers followed by flow cytometry is routinely used to interrogate T-cell responses, to characterize antigen-specific TCR repertoires and to identify immunodominant clones^3-5^. However, polyclonal repertoires are estimated to contain 10^5^-10^8^ TCRs of distinct antigen specificities^6^. Preparing libraries of properly conformed peptide/MHC-I (pMHC-I) molecules displaying an array of peptide epitopes to probe such repertoires remains a significant challenge, due to the inherent instability of empty (i.e. peptide deficient) MHC-I molecules. To circumvent the problem of unstable peptide deficient MHC-I molecules, conditional MHC class I ligands are used^7^. Such conditional ligands, bound to the MHC-I, can be cleaved by exposure to UV light^8^, or to increased temperature^9^. Upon cleavage and in the presence of a peptide of interest, a net exchange occurs where the cleaved conditional ligand dissociates and the peptide of interest associates with the MHC-I, thereby forming the desired pMHC-I complex. Such conditional ligands, however, also have limitations. The use of photo-cleavable peptides necessitates a more elaborate protein purification protocol, and may lead to increased aggregation and sample loss during the peptide exchange step. Dipeptides, which compete with the C-terminus of bound peptides to promote exchange, have been recently proposed as alternatives to catalyze peptide exchange under physiological conditions^10^.

TAPasin Binding Protein Related (TAPBPR) is a chaperone protein homologue of Tapasin involved in the quality control of pMHC-I molecules^11^. TAPBPR associates with MHC-I molecules to edit the repertoire of displayed peptides at the cell surface^12^. In a similar manner to Tapasin^13^, TAPBPR binds several MHC-I alleles *in vitro* to promote the exchange of low-for high-affinity peptides^14^. We recently performed a detailed characterization of the TAPBPR peptide exchange cycle for the murine H-2D^d^ molecule, using solution NMR^15^. This work revealed a critical role for C- and N-terminal peptide interactions with the MHC-I peptide-binding groove, which allosterically regulates TAPBPR release from the pMHC-I. Peptide binding is therefore negatively coupled to chaperone release and, conversely, the affinity of incoming peptides for the MHC-I groove is decreased by 100-fold in the presence of TAPBPR^15^ due to a widening of the MHC-I groove, as directly observed in X-ray structures of MHC-I/TAPBPR complexes^16,17^.

Here, we leverage these mechanistic insights to produce peptide-deficient MHC-I/TAPBPR complexes for two murine and one human MHC-I allotypes, independent of photo-cleavable peptide ligands. Empty MHC-I/TAPBPR complexes are stable for months, can be readily multimerized and loaded with peptides of interest in a high-throughput manner (Figure 1). Focusing on a common human allele, HLA-A*02:01, we have extended the capability of our system by incorporating our multi-modal cellular indexing technology (ECCITE-seq)^18,19^.

**Figure 1.**
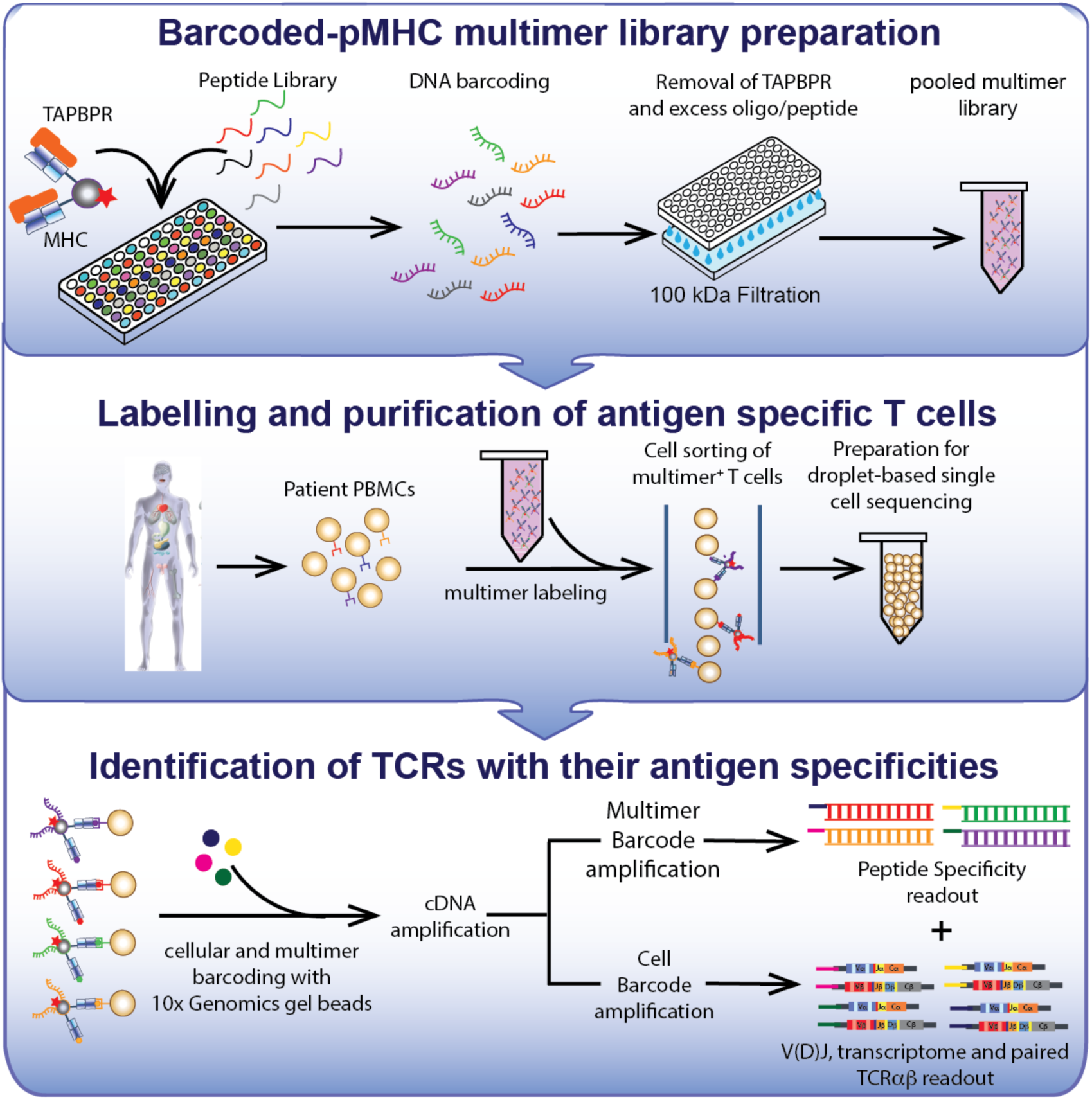
Linking peptide specificity with T-cell transcriptomes using TAPBPR-exchanged multimer libraries. Fluorophore-labelled, empty-MHC/TAPBPR multimers are loaded with peptides of interest on a 96-well plate format and individually barcoded with DNA oligos designed for 10x Genomics and Illumina compatibility. TAPBPR and excess peptide, along with free oligo, are removed by centrifugation and pooled together. A single patient sample can be stained with the pooled multimer library, and collected by fluorescence-activated cell sorting. Tetramer associated oligos and cellular mRNA from individual cells are then barcoded using 10x Genomics gel beads, followed by cDNA synthesis, library preparation and library sequencing. This workflow enables the transcriptome and paired αβ TCR sequences to be linked with pMHC specificities in a single experiment.

The resulting library of barcoded, TAPBPR exchanged tetramers can be directly applied in a multiplexed analysis of numerous antigen-specificities simultaneously, enabling the identification of TCR V(D)J sequences together with other T-cell transcriptional markers of interest in a single cell format (Figure 1).

To circumvent the need for photo-cleavable ligands, previously used to demonstrate high-affinity TAPBPR binding to empty MHC-I molecules^14^, we explored the use of destabilizing placeholder peptides. We recently described a destabilizing N-terminally truncated mutant of the P18-I10 peptide _GPGRAFVTI (gP18-I10)^15^. Here, gP18-I10 showed a high affinity for a free H-2D^d^ groove during *in vitro* refolding, but dissociated in the presence of TAPBPR to generate stable, empty H-2D^d^/TAPPBR complexes (Supplementary Fig. 1a and d). In contrast, full length P18-I10 remained captured in the groove of the H-2D^d^/TAPBPR complex (Supplementary Fig.1a-b). The previously reported 100-fold increase in the peptide off-rate for gP18-I10 relative to full-length P18-I10 peptide^15^ resulted from the loss of specific polar contacts in the H-2D^d^ A-pocket (Supplementary Fig. 2a), as further reflected in predicted IC_50_ values (24 µM gP18-I10 vs 23 nM for P18-I10). We have therefore termed this destabilizing peptide a ‘goldilocks’ peptide. Extending the same concept to a different murine MHC-I molecule, H-2L^d^, we tested an N-terminal truncation of the high-affinity gp29 epitope_PNVNIHNF (denoted gp29, IC_50_ of 16.5 µM). However, the gp29 peptide remained bound to the H-2L^d^/TAPBPR complex (Supplementary Fig. 1a and e, 2b), indicating that truncation of the extreme N-terminal residue cannot be used as a general rule to generate goldilocks peptides (Supplementary Fig. 1e). As gp29 fits the typical H-2L^d^ binding motif of xPxx[NA]xx[FLM], we explored QLSPFPFDL (QL9), a predicted low-affinity peptide (IC_50_ of 9.27 µM) with non-canonical Leu and Phe residues at position 2 and 5, respectively. Using full-length QL9 as a placeholder peptide, we could obtain empty H-2L^d^:TAPBPR complexes by further adding 10 mM Gly-Phe dipeptide (GF), which promotes peptide release by directly competing for interactions in the F-pocket of the peptide-binding groove^10^ (Supplementary Fig. 1f-g, 2b). Taken together, these results establish the principle that destabilization of peptide interactions at both ends of the groove through a range of approaches, can be used to generate peptide-deficient MHC-I/TAPBPR complexes.

To extend these findings towards a high-throughput method for the production of HLA-tetramer libraries, we focused on the common human allele HLA-A*02:01 which displays a wide range of immunodominant viral and tumor epitopes, rendering the study of HLA-A*02:01-restricted responses highly relevant^20^. Guided by the existing TAX/HLA-A*02:01 crystal structures^21^, we designed a number of mutants of the LLFGYPVYV (TAX) peptide by progressively reducing N-terminal polar contacts with MHC-I groove residues while maintaining the anchor positions 2 and 8 (x[LM]xxxxx[ILV]) (Figure 2a and Supplementary Fig. 2c). Comparison of thermal stabilities (T_m_) of HLA-A*02:01 bound to N-terminal mutants of TAX revealed a progressive reduction in T_m_ as a result of destabilization of the peptide complex upon loss of N-terminal polar contacts (Figure 2a-b). Both gTAX/HLA-A*02:01 and Ac-LLFGYPVYV/HLA-A*02:01, which gave the lowest T_m_ values at 40 °C and 46 °C, respectively, promoted the formation of peptide-deficient MHC-I/TAPBPR complexes in the presence of 10 mM dipeptide (Figure 2c, f and Supplementary Fig.3a) as shown by liquid chromatography-mass spectrometry (LC-MS) (Supplementary Fig. 3b). Empty HLA-A*02:01/TAPBPR complexes were stable at either room temperature for extended periods and could be stored at −80 °C or in lyophilized form. Incubation with the high affinity TAX peptide induced dissociation of the complex, as observed both by native gel and size exclusion chromatography (SEC) assays (Figure 2d and Supplementary Fig. 3c), with the loaded peptide detectable by LC-MS (Supplementary Fig. 3d). Accordingly, high-affinity peptides (TAX, CMVpp65 or MART-1) could be readily loaded into the empty complex, out competing fluorescently labelled TAMRA-TAX for HLA-A*02:01 binding (Figure 2e). Finally, using Bio-layer interferometry we measured the TAPBPR dissociation rate from HLA-A*02:01, whish showed a significant increase in the presence of high affinity peptides (TAX, CMVpp65 or MART-1), compared to low affinity peptides (gTAX and the irrelevant peptide gp29) (Figure 2g). This observation correlates with the respective T_m_ values of pMHC molecules refolded with different peptides (Supplementary Fig. 7c).

**Figure 2:**
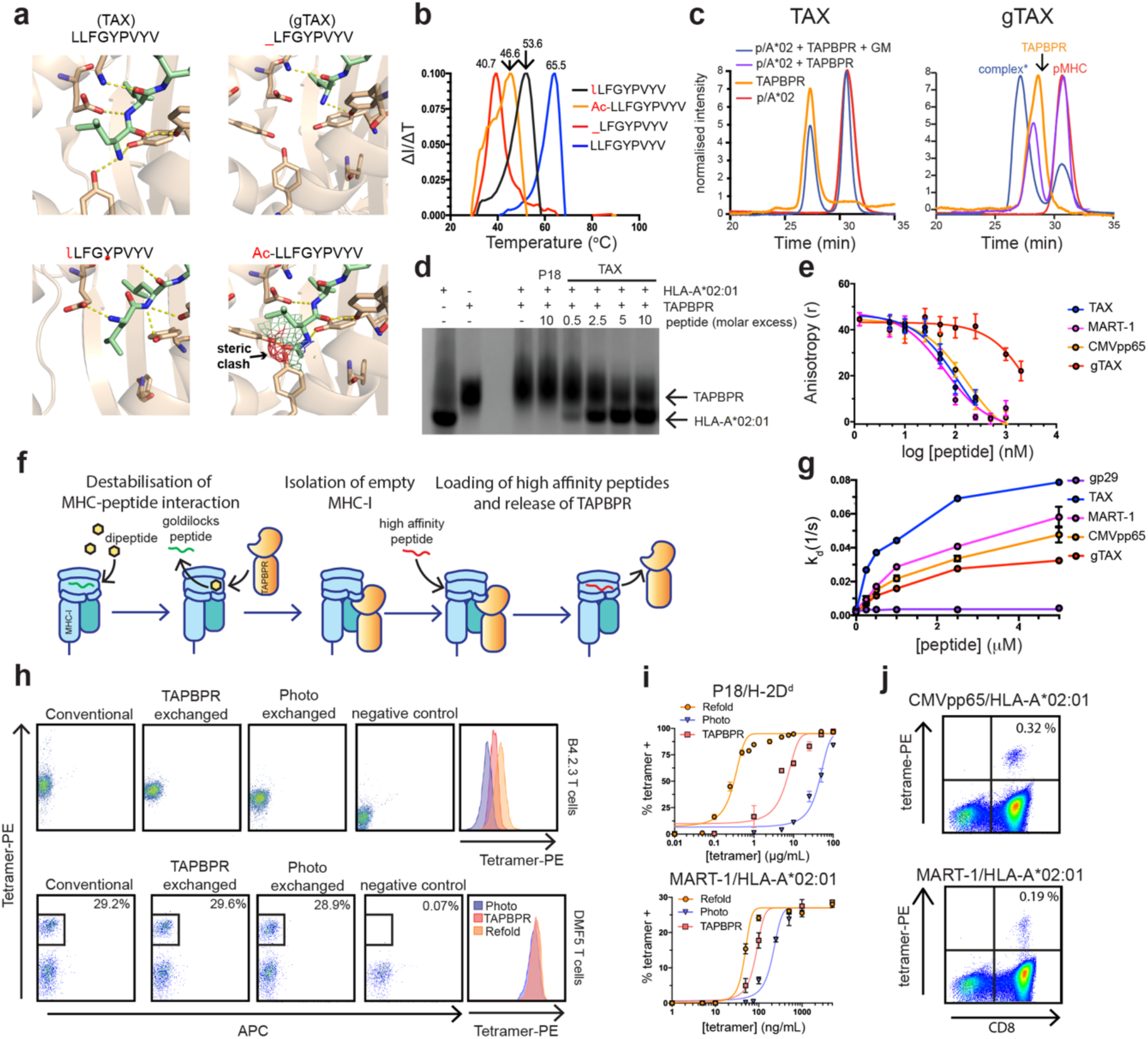
Isolation of empty MHC-I/TAPBPR complexes for high-throughput peptide exchange and tetramer library production. **(a)** Structure-based design of goldilocks peptides: comparison of polar contacts between HLA-A*02:01 and the N-terminal region of LLFGYPVYV (TAX) peptide (upper left). _LFGYPVPYV (gTAX) (upper right), Ac-LLFGYPVYV (bottom right) and ιLFGYPVYV where ι = D-Leucine (bottom left). Structures were modeled using PDB ID IDUZ^21^. **(b)** Peptide complex thermal stabilities of HLA-A*02:01 bound to TAX, lLFGYPVYV, Ac-LLFGYPVYV and gTAX. **(c)** SEC TAPBPR binding assays of TAX/HLA-A02:01 (left), gTAX/HLA-A02:01 (right). **(d)** Native gel electrophoresis of HLA-A*02:01/TAPBPR complex dissociation in the presence of relevant high affinity peptide (TAX), and irrelevant peptide (P18-I10). **(e)** Competitive binding of TAMRA-TAX to purified HLA-A*02:01/TAPBPR complexes from (c) as a function of increasing peptide concentration, measured by fluorescence polarization. **(f)** Conceptual diagram of TAPBPR-mediated capture of empty MHC-I molecules. **(g)** Bio-Layer Interferometry analysis of TAPBPR dissociation from HLA-A*02:01 in the presence of peptides of different affinity. **(h)** Representative flow cytometry analysis of tetramer staining using *in vitro* refolded (conventional), TAPBPR-exchanged and photo-exchanged tetramers. Top panels are mouse B4.2.3 TCR expressing T-cells stained with H-2D^d^:P18-I10 tetramers. Bottom panels are DMF5 TCR expressing T-cells stained with MART-1/HLA-A*02:01 tetramers. Negative controls are tetramers presenting gP18 or gTAX. **(i)** Tetramer titration of P18-I10/H-2D^d^ (top) and MART-1/HLA-A*02:01 (bottom) of conventional refolded, TAPBPR-exchanged and photo-exchanged tetramers. Data shown is representative of two independent experiments performed in duplicate where the error bars indicate standard error of the mean. **(j)** Flow cytometry of IL-2/IL-7 expanded PBMCs stained with TAPBPR-exchanged HLA-A*02:01 tetramers loaded with CMVpp65 (top plot) and MART-1 (bottom plot).

We then compared the performance of TAPBPR exchanged phycoerythrin (PE)-tetramers with PE-tetramers generated using photo-cleavable ligands^8^ or pMHC-I molecules refolded *in vitro* using standard protocols^22^. Staining of a B4.2.3 TCR transgenic T-cell line^23^ with P18-I10 /H-2D^d^ tetramers showed a two-fold enhancement in mean fluorescence intensity for TAPBPR exchanged tetramers relative to tetramers generated by UV exchange, and a decreased intensity relative to refolded pMHC-I tetramers (Figure 2h). This effect was quantified using tetramer titrations, which showed a staining saturation curve with an EC_50_ of 6.2 µg/mL compared to 42.9 µg/mL for TAPBPR and UV exchanged tetramers, respectively (Figure 2i). A similar trend was observed in MART-1/HLA-A*02:01 tetramer staining of Jurkat/MA T-cells transduced with a MART-1 specific TCR, DMF5^24^. Here, TAPBPR exchanged tetramers showed a 2.6-fold improvement in EC_50_ values relative to UV exchanged tetramers, despite staining with equivalent fluorescence intensities at saturating concentrations (Figure 2h-i). Under polyclonal sample conditions, TAPBPR exchanged tetramers were also effective at identifying low frequency CMVpp65 and MART-1 specific T-cells among cytokine expanded peripheral blood mononuclear cells (PBMCs) (Figure 2j).

The disparity between the performance of P18-I10/H-2D^d^ tetramers compared to MART-1/HLA-A*02:01 could arise from a higher binding affinity of TAPBPR for H-2D^d^ (Supplementary Fig. 1a). Since TAPBPR has been shown to promote the release of high affinity peptides from the MHC groove *in vitro*^15^, the persistence of TAPBPR following exchange may partially regenerate empty MHC-I molecules, thereby reducing the staining efficiency of the resulting tetramers. In a library format this may also lead to “scrambling” of excess peptides between MHC tetramers upon mixing, which would limit the use of molecular indices or “barcodes” to label tetramers according to their displayed peptides. Specifically, the presence of TAPBPR in the sample induced the exchange of CMVpp65 loaded on HLA-A*02:01 tetramers for free MART-1 peptide (Supplementary Fig. 4b). To prevent this, full removal of TAPBPR molecules was readily achieved by spin column dialysis, immediately following the tetramerization and peptide loading steps (Figure 1 and Supplementary Fig. 4a). The resulting pMHC-I tetramers did not capture free high affinity peptides, even when present at high (20x) molar excess, and peptide exchange between tetramers was undetectable by flow cytometry (Supplementary Fig. 4b-c).

We next sought to fine-tune TAPBPR interactions with murine MHC-I molecules previously shown to form stable complexes even when bound to high affinity peptides^15^, an effect which limits their use towards high-throughput tetramer library production. Using H-2D^d^, we explored designed mutations at the α3 domain of the heavy chain which participates in direct interactions with TAPBPR, and does not contribute to the formation of the peptide binding groove. Specifically, Met 228 is located at an edge loop of the α3 immunoglobulin fold and forms a hydrophobic contact with TAPBPR residues in the X-ray structure of the H-2D^d^/TAPBPR complex^16^ (Supplementary Fig. 5a). We hypothesized that a mutation at this position from a Met, present in H-2D^d^ and H-2L^d^, to a polar Thr residue, present in HLA-A*02:01, would lead to a reduced binding affinity of peptide-bound MHC-I molecules for TAPBPR. In contrast to WT H-2D^d^, H-2D^d^M228T did not bind TAPBPR in the presence of the high affinity P18-I10 peptide (Supplementary Fig. 5c). However, TAPBPR maintained binding by SEC to empty H-2D^d^M228T upon dissociation of the goldilocks gP18-I10 peptide, to generate a peptide-receptive complex (Supplementary Fig. 5c, d). These results highlight the requirements of a system that is amenable to large-scale tetramer library production using our approach: *i)* formation of a stable TAPBPR complex with an empty molecule and *ii)* TAPBPR dissociation upon binding of high affinity peptides to the MHC-I groove.

To test our method on a high-throughput setting we generated two tetramer libraries, one containing 29 tumor epitopes identified in neuroblastoma tissues (manuscript in preparation) and another encompassing a range of 31 viral, neoantigen and autoimmune epitopes (Supplementary Tables 1 and 2). To link pMHC specificities with TCR V(D)J sequences present in polyclonal samples, we barcoded fluorophore-labelled pMHC tetramers, prepared using TAPBPR exchange, with biotinylated DNA oligonucleotides (oligos)^25^. We used an in-house oligo design compatible with 10x Genomics gel bead oligos in the 5’ V(D)J product (Figure 1 and Supplementary Fig 6a) adding an additional modality of cellular information to our recently described ECCITE-seq method^18^. This method incorporates a cellular barcode into cDNA generated from both tetramer oligos and TCR mRNA, thus the pairing of cellular barcodes can connect TCR sequences and other mRNAs, with tetramer specificities.

Formation of stable pMHC species upon loading of each peptide was confirmed using two complementary differential scanning Fluorimetry (DSF) assays (Supplementary Fig. 7 and 8), performed on a 96-well format. After exhaustive dialysis of TAPBPR, excess barcodes and peptides, all individual peptide loading reactions were pooled into a single tetramer library sample for staining. Both final libraries prepared in this manner were further validated for: *i)* the presence of all barcodes, using bulk sequencing reactions (Supplementary Fig. 6b) and *ii*) staining of Jurkat/MA cells transduced with the DMF5 receptor specific to the MART-1 “reference” peptide included in each library. While the use of oligonucleotide barcodes leads to a reduced staining efficiency relative to non-barcoded tetramers, likely due to the lower avidity for the pMHC antigen, the observed signal intensities (10^3^-10^4^) were sufficient to distinguish the approx. 33% population of DMF5 positive cells from the negative fraction (Supplementary Fig. 6c).

To confirm that our approach has sufficient sensitivity to detect antigen-specific T-cells within a heterogeneous sample, 1% DMF5 Jurkat/MA T-cells were spiked into a sample of CD8-enriched PBMCs co-cultured with dendritic cells and stained with the library of 31 neoepitopes, including the MART-1 peptide. 100,000 tetramer positive cells were collected and 3,000 were sequenced. From this sparse sample, 256 cells with the highest number of DMF5 TCR reads were extracted and plotted together with their corresponding MART-1 tetramer reads (Figure 3a). In total, we recovered 85 cells with ≥10 MART-1 tetramer counts, 76 of which showed ≥ 10 DMF5 TCR reads (considered DMF5 positive) giving an approximate false positive rate of 10.6% (Figure 3a). Conversely, 6 DMF5 TCR positive cells showed ≤ 10 MART-1 tetramer counts, resulting in a false negative rate of 7.9%. The low number of cells with significant tetramer barcode reads could be a function of tetramer avidity, exacerbated by high dilutions through the 10x Genomics system prior to cDNA preparation. Overall, a clear positive relationship between MART-1 counts and DMF5 TCR reads was observed, suggesting that our tetramer libraries can be used to identify sparse populations of antigen-specific T-cells within a heterogeneous sample.

**Figure 3:**
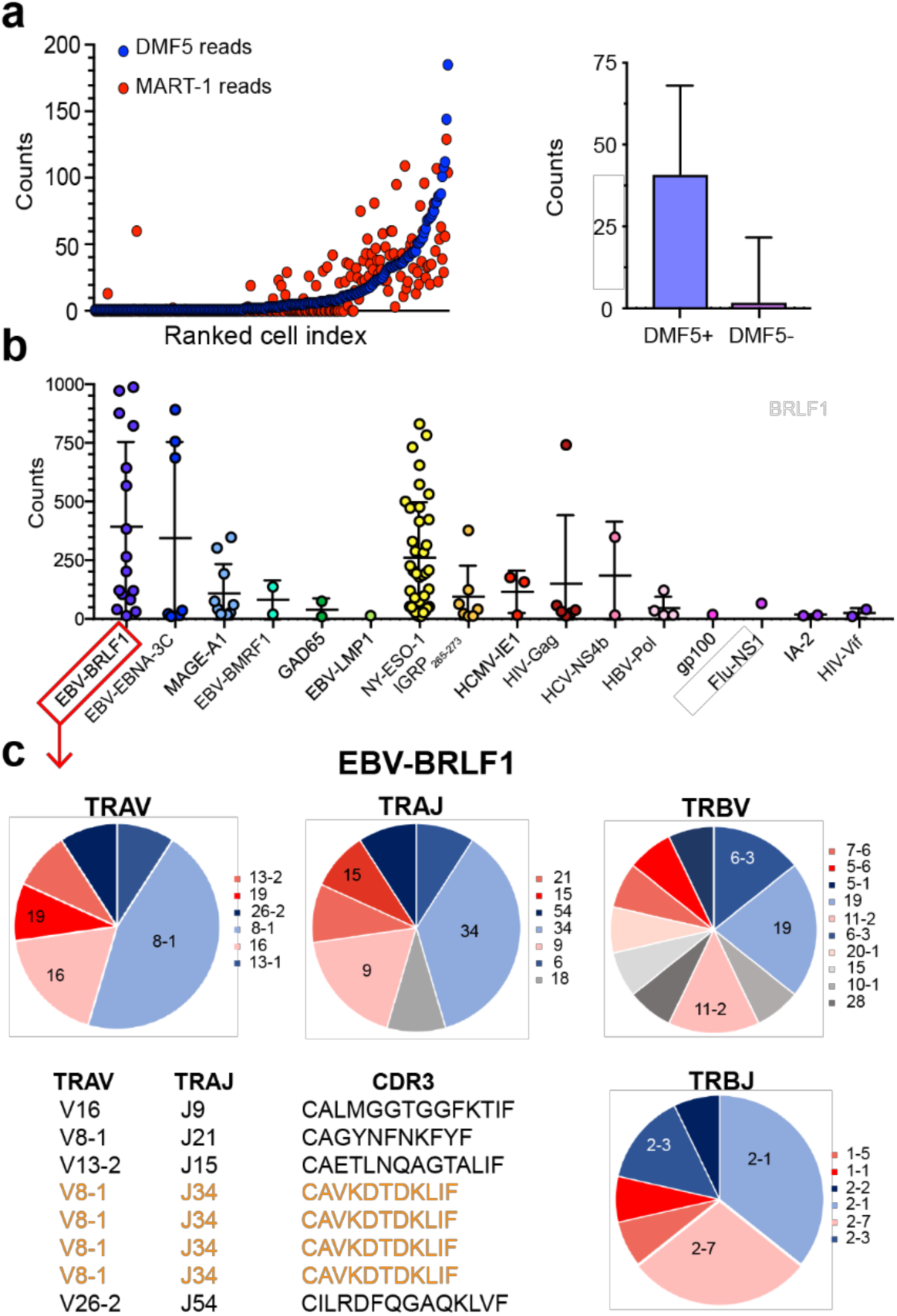
Identification of paired αβ TCR sequences with their pHLA-A*02:01 antigen specificities from polyclonal T-cell samples. **(a)** Recovery of MART-1 tetramer barcodes on DMF5 cells from PBMC-DC co-cultures spiked with 1% DMF5 cells. Number of MART-1 tetramer counts and number of DMF5 counts (both plotted on the y-axis) per cell analyzed (x-axis), where n = 256. Cells are ordered based on DMF5 read counts. Bar graph displays the mean MART-1 tetramer reads from DMF5+ (>10 DMF5 reads n = 76) and DMF5-(≤ 10 DMF5 reads, n = 927) cells. Error bars indicate 1 standard deviation from the mean. **(b)** Distribution of antigen specificities identified from tetramer+/CD8+ T-cells from human splenocytes and the number of tetramer-barcode read per cell. Each dot represents a single cell, n =102 in total. **(c)** V(D)J usage of cells identified as specific for the EBV-BRLF1 antigen (YVLDHLIVV). All TRAVJ (n = 11) and TRBVJ (n = 14) chains identified are represented. TCR alpha chains identified for EBV-BRLF1 specific T-cells. Repeated BRLF-1 TRA chains are highlighted in orange. Gating strategies used for sorting tetramer+ cells are outlined in Supplementary Fig. 9.

Finally, we used our methodology to probe distinct TCR specificities present in a polyclonal repertoire of T-cells. A total of 2 × 10^6^ CD8^+^ T-cells isolated from the spleen of an EBV-positive donor were stained using our barcoded tetramer library consisting of 34 viral, autoimmune and tumor epitopes (Supplementary Table 2). After sorting, 3,000 tetramer positive cells were loaded on the 10x platform for single-cell sequencing. ECCITE-seq analysis retrieved 1,722 cell barcodes, the majority of which were associated with < 2 tetramer reads, giving a low apparent background. 110 cells were further identified as tetramer-enriched, defined as cells with >10 tetramer reads for at least one peptide specificity. 27 tetramer-enriched cells bound to >5 different peptide epitopes, 5 of which showed no bias towards a particular epitope and were excluded from subsequent analysis. Among the final set of 102 tetramer-enriched cells, we identified at total of 16 distinct epitope specificities, with an average of 200 reads per cell for each dominant tetramer (Figure 3b). Specifically, a large fraction of tetramer-enriched cells had high reactivity for the NY-ESO-1 epitope (37 cells), followed by EBV-BRLF1 (16 cells), MAGE-A1 (10 cells) and IGRP_265-273_ (7 cells). Towards validating the significance of our results, we focused on barcodes corresponding to the EBV-BRLF1 epitope whose TCR repertoire has been previously characterized using *ad hoc* tetramers prepared using conventional refolding protocols^26^. Inspection of V(D)J TCR sequence reads from tetramer-enriched cells identified a clear bias towards the usage of TRAV8-1 (50%) with TRAJ34 (50%), TRBV19 (21%), TRBJ2-1 (36%) and TRBJ2-7 (29%), a finding consistent with published reports on EBV-BRLF1 specific repertoires^27^. Whereas the β-chain CDR3 sequences were more variable, analysis of TRA CDR3 sequences further identified a known public alpha chain CDR3 (CAVKDTDKLIF) which was previously linked to a functional TCR, specific for the EBV-BRLF1 epitope^26^. This sequence was observed in half of EBV-BRLF1 tetramer-enriched cells, and was not detected in cells enriched for any other tetramer (Figure 3e). Taken together, these results corroborate our initial findings using the spiked DMF5/PBMC sample (Figure 3a), and further suggest that our barcoded pMHC libraries prepared using TAPBPR exchange are of sufficient quality and staining efficiency to identify antigen-specific TCRs present within polyclonal repertoires.

We have outlined a robust method to isolate stable, empty MHC-I molecules at milligram quantities, that can be readily loaded with peptides of choice in a high-throughput manner. Rather than relying on chemically synthesized conditional ligands, our approach employs a molecular chaperone, TAPBPR, in an analogous manner to the antigen processing pathway used by cells to load MHC-I molecules with immunodominant peptides^1^. In combination with a simple indexing design that is compatible with the 10x Genomics system, our approach can efficiently link paired TCR V(D)J sequences and other transcription markers to their pMHC specificities in a polyclonal sample setting. Our peptide exchange and barcode sequencing workflow has no theoretical upper limit with respect to library size, other than the cost of peptide and oligo synthesis, which renders it suitable for the simultaneous analysis of hundreds of epitope specificities in future experiments. In addition to expediting tetramer library preparation and the identification of novel TCR specificities, our method can be readily extended to include the analysis of complete transcriptomes^18,19^, thereby providing a toehold for functional studies of TCR:pMHC recognition.

## AUTHOR CONTRIBUTIONS

N.G.S. conceived and designed the project. S.A.O. produced and characterized murine MHC-I/TAPBPR complexes, analyzed V(D)J sequencing data, and together with D.M. produced recombinant TAPBPR protein. J.S.T. and D.M. produced and characterized HLA-A*02:01/TAPBPR complexes, J.S.T. performed Fluorescence anisotropy and bio-layer interferometry assays and made tetramer libraries. N.G.S., M.Y. and P.S. conceived and designed the linking of binding specificity to TCR sequences with oligo barcoded tetramers, S.H. and P. S. performed sequencing experiments and curated sequence data. M.Y., S.N., A.J., M.B. and J.M.M. designed experiments using polyclonal cell samples, provided all samples and performed flow cytometry and sorting experiments. S.A.O. and N.G.S. wrote the manuscript, with feedback from all authors.

## ACKNOWLEDGEMENTS

We would like to thank Brian Fritz (10x Genomics), Andrew McShan (UCSC), Marlon Stoeckius (NYGC) and Erik Procko (UIUC) for helpful discussions throughout the course of this study. We are grateful to Kannan Natarajan and David Margulies (NIH) for providing the TAPBPR S2 and B4.2.3 TCR mammalian cell lines, Alejandro Rodriguez and Jevgenij Raskatov for synthesis of Ac-LLFGYPVYV and lLFGYPVYV peptides, Brett Thurlow and Charles Heffern (nanoTemper), for recording and analyzing nanoDSF data. Work at NYGC was supported by grants to P.S. from the NIH (R21HG009748) and the Chan Zuckerberg Initiative (HCA-A-1704-01895). This work was also supported by grants from the National Institutes of Health to N.G.S. (R35GM125034, R01AI143997), to M.R.B. (UC4 DK112217), and to J.M.M. (R35CA220500, U54CA232568).

## ONLINE METHODS

### Peptides

All peptide sequences are given as standard single letter code. Peptides used for MHC refoldings and production of the neoepitope library were purchased from Genscript at 98% purity or as pepsets from Mimotopes as crude peptides and dissolved in 8.25% Acetonitrile, 25% DMSO, and 66.75% H_2_O. Peptides containing modifications: TAMRA-TAX and GILGFVFXL (where X = 3-amino-3-(2-nitrophenyl)-propionic acid) were purchased from Biopeptik at 98% purity. lLFGYPVYV and Ac-LLFGYPVYV were synthesized in house using standard FMOC chemistry. Peptide binding affinities were predicted using netMHCpan 4.0^28^.

### *In vitro* refolding of pMHC molecules

Plasmid DNA encoding the luminal domain of class I MHC (MHC-I) heavy chains H-2D^d^, HLA-A*02:01, H-2L^d^ and human β_2_-microglobulin (hβ_2_m,) were provided by the tetramer facility (Emory University), and transformed into *Escherichia coli* BL21(*DE3)* (Novagen). MHC-I proteins were expressed in Luria-Broth media, and inclusion bodies (IBs) were purified as previously described^22^. *In vitro* refolding of pMHC-I molecules was performed by slowly diluting a 200 mg mixture of MHC-I and hβ2m at a 1:3 molar ratio over 24 hours in refolding buffer (0.4 M L-Arginine, 100 mM Tris pH 8, 2 mM EDTA, 4.9 mM reduced glutathione, 0.57 mM oxidized glutathione) containing 10 mg of synthetic peptide purchased from Genscript at 98% purity at 4 °C. H-2D^d^ heavy chain was refolded with RGPGRAFVTI (P18-I10) derived from HIV gp120^29^ or _GPGRAFVTI (gP18-I10). H-2L^d^ was refolded with _PNVNIHNF (gp29) or QLSPFPFDL (QL9) derived from oxo-2-gluterate dehydrogenase. HLA-A*02:01 was refolded with variants of LLFGYPVYV (TAX) derived from HTLV-1 including _LFGYPVYV (gTAX), N-terminally acetylated TAX (Ac-LLFGYPVYV), lLFGYPVYV where the first residue is a D-leucine or with ELAGIGILTV (MART-1) derived from Melan-A. Refolds were allowed to proceed for 96 h followed by size exclusion chromatography (SEC) using a HiLoad 16/600 Superdex 75 column (150 mM NaCl, 25mM Tris pH 8) at a flow rate of 1 mL/min, followed by anion exchange chromatography on a mono Q 5/50 GL column at 1 mL/min using a 40 minute 0-100% gradient of buffer A (50 mM NaCl, 25mM Tris pH 8) and buffer B (1M NaCl, 25mM Tris pH 8). Typical protein yields from a 1L refold were 5 to 10 mg of purified pMHC-I.

### Recombinant TAPBPR expression and purification

The luminal domain of TAPBPR was expressed using a stable *Drosophila* S2 cell line (Dr Kannan Natarajan, National Institutes of Health) induced with 1 mM CuSO_4_ for 4 days and purified as described previously^16^. Briefly, His_6_-tagged TAPBPR was captured from the supernatant by affinity chromatography using high-density metal affinity agarose resin (ABT, Madrid). Eluted TAPBPR was further purified by size exclusion using a Superdex 200 10/300 increase column at a flow rate of 0.5 mL/min in 100 mM NaCl and 20 mM sodium phosphate pH 7.2.

### Size exclusion chromatography of MHC-I/TAPBPR complexes

SEC analysis of MHC-I/TAPBPR interaction was performed by incubating 40 µM purified pMHC-I molecules with purified TAPBPR at a 1:1 molar ratio in 100 mM NaCl, 20 mM sodium phosphate pH 7.2 for 1 h at room temperature. Complexes were resolved on an Superdex 200 10/300 increase column (GE healthcare) at a flow rate of 0.5 mL/min in 100 mM NaCl and 20 mM sodium phosphate pH 7.2 at room temperature. MHC-I/TAPBPR complexes eluted at 26.5 min. In the case of H-2L^d^ and HLA-A*02:01, 10 mM GF and GM were added respectively both to the initial incubation and to the running buffer during chromatography.

### LC-MS

Peptide occupancy of SEC purified MHC-I was determined by HPLC separation on a Higgins PROTO300 C4 column (5 µm, 100 mm × 21 mm) followed by electrospray ionisation performed on a Thermo Finnigan LC/MS/MS (LQT) instrument. Peptides were identified by extracting expected *m/z* ions from the chromatogram and deconvoluting the resulting spectrum in MagTran.

### Preparation of photo-exchanged pMHC-I

H-2D^d^ refolded with RGPGRAFJ*TI (photo-P18-I10) and HLA-A*02:01 refolded with GILGFVFJ*L, where J* is the photo-cleavable residue 3-amino-3-(2-nitrophenyl)-propionic acid, were UV-irradiated at 365 nm for 1 h in the presence of 20-fold molar excess peptide at room temperature. Reactions were iced for 1 h then centrifugated at 14, 000 rpm for 10 min to remove aggregates. Photo-exchanged pMHC-I was then used for DSF analysis or tetramer preparation.

### Differential scanning Fluorimetry (DSF)

To measure thermal stability of pMHC-I molecules^30^, 2.5 µM of protein was mixed with 10 × Sypro Orange dye in matched buffers (20 mM sodium phosphate pH 7.2, 100 mM NaCl) in MicroAmp Fast 96 well plates (Applied Biosystems) at a final volume of 50 µL. DSF was performed using an Applied Biosystems ViiA qPCR machine with excitation and emission wavelengths at 470 nm and 569 nm respectively. Thermal stability was measured by increasing the temperature from 25 °C to 95 °C at a scan rate of 1 °C/min. Melting temperatures (T_m_) were calculated in GraphPad Prism 7 by plotting the first derivative of each melt curve and taking the peak as the T_m_ (Supplementary Fig.7a). Determination of T_m_ values of TAPBPR exchanged molecules additionally required subtraction of the TAPBPR melt curve from the curve obtained for the complex, then calculating the first derivative. This procedure, on average, enhanced the T_m_ values calculated for TAPBPR exchanged pMHC-I molecules by 1.5 °C, compared to refolded and photo-exchanged pMHC-I molecules. All samples were analyzed in duplicate and the error is represented as the standard deviation of the duplicates analyzed independently.

### Bio-Layer Interferometry

In each experiment, HIS1K biosensor tips (ForteBio) were first baselined in a buffer of 20 mM sodium phosphate pH 7.2, 100 mM NaCl and then coated with 6 µg/mL of HIS-Tagged TAPBPR in a matched buffer until the response was between 0.3 nm and 0.4 nm for each tip. TAPBPR coated biosensor tips were then dipped into matched buffer supplemented with 0.02% TWEEN-20 and 0.5 mg/mL BSA for 6 minutes to block non-specific interaction and as secondary baseline step. Subsequent steps were performed in the secondary baseline buffer (20 mM sodium phosphate pH 7.2, 100 mM NaCl, 0.02% TWEEN-20, 0.5% BSA). Biosensor tips were then dipped into buffer containing 10 µM HLA-A*02:01/g-TAX and 10 mM GM dipeptide to facilitate peptide deficient MHC/TAPBPR formation for 10 minutes. After peptide deficient MHC/TAPBPR formation on the biosensor tips, they were dipped into buffer supplemented with 0-5 µM of the indicated peptides for 14 minutes to facilitate pMHC dissociation from TAPBPR. Data was processed after subtraction of the reference sensor tip data set which was coated with TAPBPR in buffer, Y-axis alignment to the secondary baseline, and an interstep correction alignment to dissociation. All data was locally fit for association and dissociation with a 2:1 (heterogenous) binding model. Goodness of fit was determined according to R^2^ values of above 0.99. All experiments were performed using an Octet Red 96e system and analyzed with the Octet data analysis HT v.11.1 software.

### Nano Differential Scanning Fluorimetry (nanoDSF)

In a volume of 20 µl, 1 µM peptide deficient HLA-A*02:01/TAPBPR was incubated with 20 µM of various peptides (Supplementary Table 1) in a buffer of 20 mM sodium phosphate pH 7.2, 100 mM NaCl, 0.02% TWEEN-20 for at least one hour. 10 uL of each sample was loaded on the Prometheus NT.48 instrument (NanoTemper) using the high sensitivity capillaries. NanoDSF measurements are performed using a temperature ramp rate of 1°C/min from 25 °C to 95 °C and an LED intensity of 20%. Data was analyzed using the PR Control software (NanoTemper). Melting temperatures (T_m_ values) correspond to the inflection points of the first derivative of the 330/350 nm fluorescence ratio. Experiments were performed in duplicates and the shaded areas (Supplementary fig. 7b) are representative of the error.

### Native gel electrophoresis

Peptide-deficient MHC-I/TAPBPR complexes were incubated with the indicated molar ratio of relevant (TAX) or irrelevant (P18-I10) peptide for 1 h at room temperature. Samples were run at 90 V on 8 % polyacrylamide gels in 25 mM TRIS pH 8.8, 192 mM glycine, at 4°C for 4.5 hours and developed using InstantBlue (Expedeon).

### Fluorescence anisotropy

Fluorescence anisotropy was performed using TAX peptide labeled with TAMRA dye (K_TAMRA_LFGYPVYV) (herein called TAMRA-TAX), as described previously for the H-2D^d^ system^15^. Briefly, 50 nM of peptide-deficient HLA-A*02:01/TAPBPR complex in 100 mM NaCl, 20 mM sodium phosphate and 0.05 % (v) tween-20, were incubated with 0.75 nM TAMRA-TAX and graded concentrations of MART-1, CMVpp65 or unlabeled TAX peptide in total volumes of 100 µL in black 96 well assay plate (Costar 3915) for 2 hours at room temperature while Fluorescence anisotropy (r) was recorded. Fluorescence anisotropy (FA) was recorded on a Perkin Elmer Envision 2103 with an excitation filter of *λ*_ex_ = 531 nm and an emission filter of *λ*_em_ = 595 nm. FA were normalized against TAMRA-TAX alone. Measurements were recorded every 30 seconds and data points represented are an average of FA values acquired following 105 minutes of incubation. All experiments are representative of at least 3 individual experiments run in triplicates. Data points were plotted and fit using a sigmoidal response curve in GraphPad Prism 7.

### Preparation of barcoded peptide exchanged tetramers

Purified, *BirA*-tagged pMHC-I molecules were biotinylated using the Bulk *BirA: BirA* biotin-protein ligase bulk reaction kit (Avidity), according to the manufacturer’s instructions. Biotinylated pMHC-I was buffer exchanged into PBS pH 7.4 using a PD-10 desalting column. Biotinylation was confirmed by SDS-PAGE in the presence of excess streptavidin and by LC-MS. Biotinylated peptide-deficient MHC-I/TAPBPR complexes were generated and purified by SEC as described above. Empty-MHC-I/TAPBPR complex eluting at 26.5 min, was collected and confirmed to contain both MHC-I and TAPBPR by SDS-PAGE. Purified complexes were confirmed peptide-deficient by LC-MS and stored at −80 °C. Tetramerization of empty-MHC-I/TAPBPR was performed by adding a 2:1 molar ratio of biotinylated MHC-I/TAPBPR to streptavidin-PE or streptavidin-APC (Prozyme) in five additions over 1 h on ice. Tetramerized empty-MHC-I/TAPBPR complexes were then aliquoted into 96 well plates at 2 μg of total MHC-I per well. For tetramer barcoding, custom biotinylated DNA oligos (IDT) were diluted to a concentration of 8.5 µM prior to use. Each tetramer was barcoded by adding 2:1 molar equivalent of DNA-barcodes relative to streptavidin and incubated for 1 hr, at 4 °C. Peptide-deficient barcoded-MHC-I/TAPBPR tetramers were then exchanged with peptides of interest by adding a 20-molar excess of peptide to each well and incubating for 1 hour. Additionally, 8 molar excess biotin (to block any free streptavidin sites) was added and incubated for a further 1 h at room temperature. After exchange, tetramers were transferred to 100 kDa spin columns (Amicon) and washed with 1000 volumes of PBS to remove TAPBPR and excess peptide and barcodes. After washing, exchanged tetramers were pooled and stored at 4 °C for up to 3 weeks.

### Cell Culture

58 α^−^β^−^ T-cells expressing the B4.2.3 TCR, which recognizes P18-I10 bound to H-2D^d^, were obtained from Dr. Kannan Natarajan (NIH). TCR β-chain deficient Jurkat-MA T-cells expressing the DMF5 TCR, which recognizes Melan-A epitope MART-1 bound to HLA-A*02:01. Cells were grown in DMEM supplemented with 10 % FBS, 25 mM HEPES pH 7, 2 µM β-mercaptoethanol, 2 mM L-glutamine, 100 U/mL penicillin/streptomycin and 1 × non-essential amino acids. Cells were maintained in exponential phase in a humidified incubator at 37 °C with 5 % CO_2_. Splenocytes from HLA-A*02:01+ organ donors were obtained through the Human Pancreas Analysis Program (http://hpap.pmacs.upenn.edu/) (University of Pennsylvania) after informed consent by each donor’s legal representative.

For cytokine expansion, PBMCs from healthy HLA-A*02:01 matched donors (Children’s Hospital Philadelphia) were cultured in X-VIVO 15 (Lonza) supplemented with 10 % human serum albumin, 25 mM HEPES pH 7, 2 µM β-mercaptoethanol, 2 mM L-glutamine, 100 U/mL penicillin/streptomycin, 1x non-essential amino acids, 1000 U/mL recombinant human IL-2, 10 ng/mL recombinant human IL-7 and 10 ng/mL recombinant human IL-15 for 2 weeks prior to analysis.

### Generation of DMF5 T-cell line

Retrovirus for transduction of Jurkat/MA^31^ and primary CD8 T-cells was produced using Platinum-A retroviral packaging cell line. DMF5 cassette was assembled using previously described CDR3 sequences and V(D)J family genes^32^, codon optimized, synthesized, and cloned into pMP71 retroviral vector^33^. Jurkat/MA cells were plated in 6-well plates at 7 × 10^5^ cells/well and transfected with 2.5 mg of retroviral vector pMP71 using Lipofectamine 3000 (Life Technologies, Invitrogen). After 24 hours, medium was replaced with IMDM-10 % FBS or AIM-V-10 % FBS. Supernatants were harvested and filtered with 0.2 mM filters after 24 hours incubation and transferred to Jurkat/MA cells in 6-well plates pre-treated with 1 mL well/Retronectin (20 mg/mL in PBS, Takara Bio. Inc.,) at 1 × 10^6^ cells/well and spinoculated with 2 mL of retroviral supernatant at 800 g for 30 min at RT. After 24 hours, cells were washed and PBS, and cultured in IMDM-10% FBS. Jurkat/MA cells were stained with MART-1 dextramer and sorted for dextramer (Immudex) positive cells.

### PBMC/DC co-culture

Normal donor monocytes were plated on day 1 in 6-well plates at 5 × 10^6^/ well in RPMI-10 FBS supplemented with 10 ng/ml IL-4 (Peprotech) and 800 IU/ml GM-CSF (Peprotech) and incubated at 37 °C overnight. On day 2, fresh media supplemented with 10 ng/ml IL-4 and 1600 IU/ml GM-CSF was added to the monocytes and incubated at 37°C for another 48 hours. On day 4, non-adherent cells were removed and immature dendritic cells washed and pulsed with 5 uM peptide in AIM-V-10 % FBS supplemented with 10 ng/ml IL-4, 800 IU/ml GM-CSF, 10 ng/ml LPS (Sigma-Aldrich), and 100 IU/ml IFN-γ (Peprotech) at 37°C overnight. Day 1 was repeated on days 4 and 8 to generate dendritic cells for the second and third stimulations on days 8 and 12, respectively. On day 5, normal donor-matched CD8+ T-cells were co-cultured with the pulsed dendritic cells in AIM-V-10 % FBS. Day 5 protocol was repeated on day 8 and day 12 using dendritic cells generated on days 4 and 8 for the second and third stimulation, respectively.

### Flow cytometry

Tetramer analysis was carried out by staining 2 × 10^5^ cells with anti-CD8*α* mAb (BD Biosciences) and 1 μg/mL HLA-A02:01/MART-1 tetramer or 50 μg/mL H-2D^d^/P18-I10 tetramer for 1 h on ice, followed by two washes with 30 volumes of FACS buffer (PBS, 1% BSA, 2 mM EDTA). All flow cytometric analysis was performed using a BD LSR II instrument equipped with FACSDiva software (BD Biosciences). For cell sorting experiments, cryopreserved human splenocytes were thawed and rested in RPMI media (10 % FBS, 1 % L-glutamine, 1 % Pencillin/Streptomycin). CD8^+^ T-cells were enriched by negative selection using magnetic beads according to the manufacturer’s protocol (STEMCELL Technologies). Cells were then treated with dasatinib (50 nM, Sigma-Aldrich) for 30 minutes prior to staining. Afterward, 50 μL each of PE and APC versions of the tetramer library were added (final amount was 0.5 μg pMHC per tetramer) were added for 15 minutes at room temperature. Cells were stained with anti-CCR7–APC-Cy7 (G043H7, BioLegend) for 10 minutes at 37 °C; with LIVE/DEAD Fixable Aqua (Thermo Fisher Scientific) for 10 minutes at room temperature; with an antibody cocktail containing: anti-CD14–BV510 (M5E2, BioLegend), anti-CD19– BV510 (HIB19, BioLegend), anti-CD3-APC-R700 (UCHT1, BD Biosciences), anti-CD4–PE-Cy5.5 (S3.5, Thermo Fisher Scientific), anti-CD8–BV605 (RPA-T8, BioLegend), and anti-CD45RA PE-CF594 (HI100, BD Biosciences) for 20 minutes at room temperature. Cells were washed and resuspended in BD pre-sort buffer (BD Biosciences). Cell sorting was performed on a FACS Aria FUSION (BD Biosciences)

Live cells were gated based on forward and side scatter profiles and data was analyzed using FlowJo software (FlowJo, LLC). For EC_50_ determination, tetramer concentrations were calculated based on total amount of pMHC-I at the time of exchange. Titrations were performed on the appropriate cell line in duplicate in two independent experiments. The percentage of tetramer+ T-cells was measured relative to the staining achieved at the highest concentration tested within each experiment. EC_50_ values were calculated by fitting a Boltzmann sigmoidal function to the data with the lower constraint set to 0 and upper constraints set to 95 for B4.2.3 and 28 for DMF5 in GraphPad Prism 7.

### ECCITE-seq

Post sorting, samples were prepped for the 10X Genomics 5P V(D)J kit workflow, and processed according to the ECCITE-seq protocol^18^, with these modifications:

1. For cDNA amplification, 1uL of 0.2uM tetramer additive (**GTCTCGTGGGCTCGGAGATG)** was spiked into the reaction.
2. Post cDNA PCR, a 0.6x SPRI cleanup was performed, resulting in the larger cDNA fragments being retained on the beads, and the tetramer tags in the supernatant. After separation of the two fractions and elution from the beads, a portion of the cDNA was used to perform TCR alpha/beta amplification and library prep, as described in the 10X genomics protocol.
3. A separate portion of the cDNA elution was used to perform a DMF5 receptor specific enrichment, using a hemi-nested PCR strategy akin to that used for the TCRα/β enrichment. All PCRs were performed using 2X KAPA Hifi Master Mix. Primers for PCRs: PCR - DMF5_PCR1 (GAAATTCACGGCGCACAGG) with SI-PCR primer (10x). PCR2 - DMF5_PCR2 (CCTTGGCACCCGAGAATTCCAGCTTGGCTGGCTGTCTCTGATC) and P5_generic (AATGATACGGCGACCACCGAGATCTACAC). PCR3 (to add P7 end and sample index) - RPI-x primer (“x” nucleotides comprise a user-defined index) CAAGCAGAAGACGGCATACGAGATxxxxxxxxGTGACTGGAGTTCCTTGGCA CCCGAGAATTCCA and P5_generic.
4. The supernatant of the 0.6X SPRI purification in step 2 above was purified with 2 rounds of 2x SPRI. First, 1.4x SPRI was added to the supernatant to bring up the volume to 2x, followed by two rounds of 80% ethanol washes. After eluting in water, an additional 2x SPRI cleanup was performed. Post second cleanup, the tetramer tags were converted to a sequenceable library by PCR with SI-PCR and N7XX (“x” nucleotides comprise a user-defined index) CAAGCAGAAGACGGCATACGAGATxxxxxxxxGTCTCGTGGGCTCGG.

### Sequencing and Analysis

Individual tetramers were pooled in one library sample prior to sequencing. Samples were sequenced on a Miseq using a v2 300 cycle kit (151 cycles R1, 8 cycles I1, 151 cycles read 2). Post sequencing, TCR fastq files were pooled together for each sample, then analyzed using cellranger vdj 3.0.0 against the GRCh38 reference genome (v2.0.0, as provided by the 10X website). To identify the DMF5 receptor, we used CITE-seq-Count version 1.4.1 to search for the DMF5 specific tag, using default parameters (hamming distance set to 5). For tetramers, we used CITE-seq-Count version 1.4.1 using all default parameters, with the exception of hamming distance set to 1, and a whitelist to search for only cells with TCR found by 10X.

### TCR Repertoire Analysis

All analysis was performed using the PE-tetramer barcodes alone. Cells with ≥10 tetramer reads were clustered into pMHC specificity groups based on the tetramer barcode read. Cells with multiple tetramers with >10 reads were clustered based on the most frequent tetramer read (≥ 50 % of total tetramer reads for that T-cell). All TCR sequences identified (partial or complete) were used in global VJ usage analysis. In cases where multiple TCRs were read, only TCR sequences with the highest true reads were used (generally representative of ≥ 90 % of TCR reads for that T-cell). Known receptors and CDRs were queried and identified using VDJdb and literature searches.

