## Supplementary Information for "High Throughput pMHC-I Tetramer Library Production Using Chaperone Mediated Peptide Exchange"

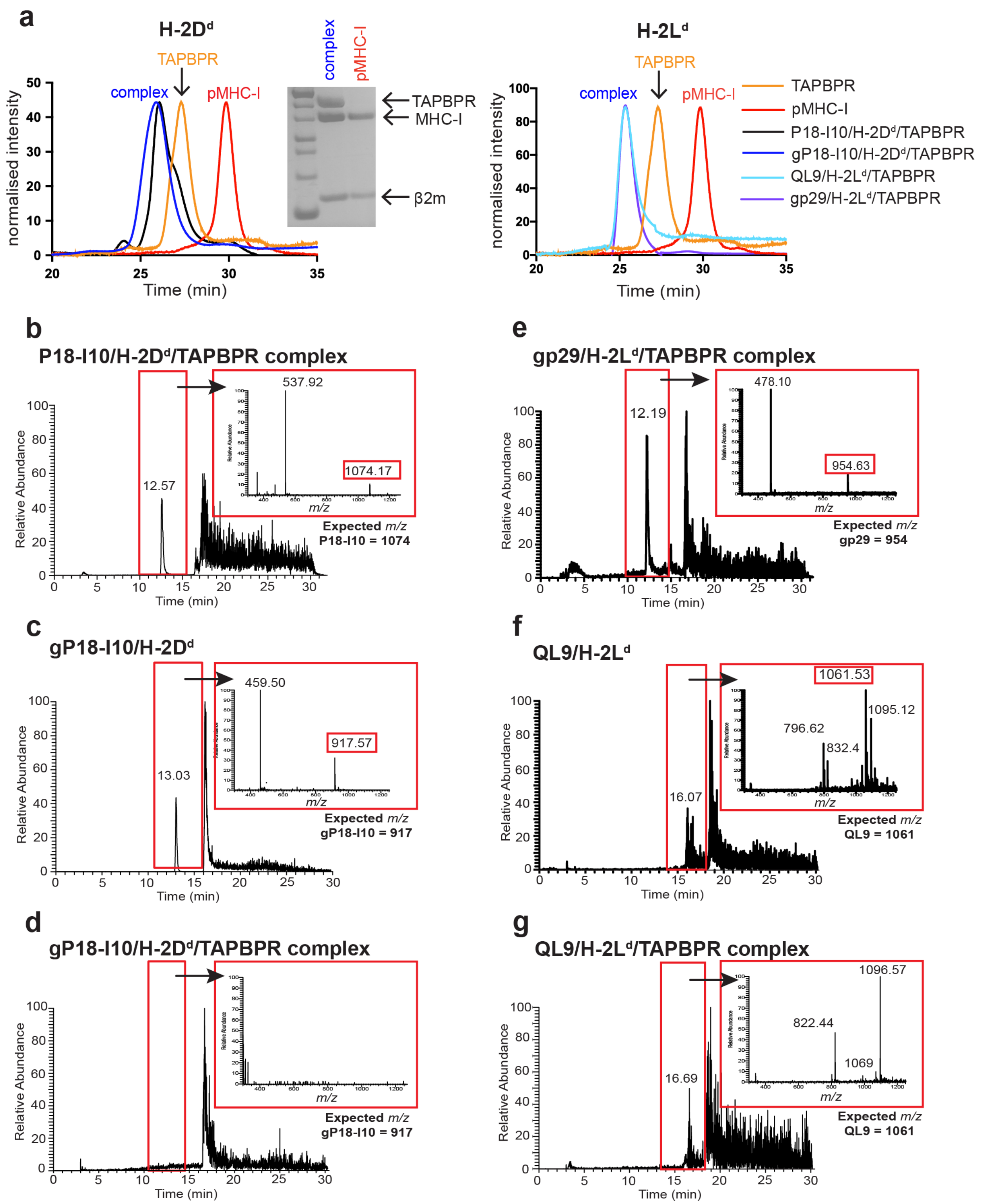

a
H-2Dd
TAPBPR
complex
pMHC-I
normalised intensity
Time (min)
TAPBPR
MHC-I
β2m
H-2Ld
TAPBPR
complex
pMHC-I
normalised intensity
Time (min)
TAPBPR
pMHC-I
P18-I10/H-2Dd/TAPBPR
gP18-I10/H-2Dd/TAPBPR
QL9/H-2Ld/TAPBPR
gp29/H-2Ld/TAPBPR
b
P18-I10/H-2Dd/TAPBPR complex
Relative Abundance
Time (min)
12.57
537.92
1074.17
m/z
Expected m/z
P18-I10 = 1074
c
gP18-I10/H-2Dd
Relative Abundance
Time (min)
13.03
459.50
917.57
m/z
Expected m/z
gP18-I10 = 917
d
gP18-I10/H-2Dd/TAPBPR complex
Relative Abundance
Time (min)
m/z
Expected m/z
gP18-I10 = 917
e
gp29/H-2Ld/TAPBPR complex
Relative Abundance
Time (min)
12.19
478.10
954.63
m/z
Expected m/z
gp29 = 954
f
QL9/H-2Ld
Relative Abundance
Time (min)
16.07
1061.53
1095.12
796.62
832.4
m/z
Expected m/z
QL9 = 1061
g
QL9/H-2Ld/TAPBPR complex
Relative Abundance
Time (min)
16.69
1096.57
822.44
1069
m/z
Expected m/z
QL9 = 1061

**Supplementary Figure 1. Destabilizing "goldilocks" peptides promote isolation of empty murine MHC-I/TAPBPR complexes. (a)** Size exclusion chromatography (SEC) elution profiles of H-2D$^d$ bound to RGPGRAFVTI (P18-I10) or GPGRAFVTI (gP18-I10) (left panel) and H-2L$^d$ bound to _PNVNIHNF (gp29) or QLSPFPFDL (QL9) (right panel) in the presence or absence of TAPBPR. SEC of H-2L$^d$ molecules shown was performed in the presence of 10 mM GF dipeptide. Inset shows SDS-PAGE analysis of elution fractions at 26 min / 18 mL (pMHC-I/TAPBPR complex) and 30 min / 15 mL (pMHC-I alone). **(b-g)** Analysis of MHC-I peptide occupancy by LC-MS. Chromatograms shown are filtered to only display peaks containing *m/z* ions of interest. Insets show MS analysis of the region indicated by the red box. **(b)** P18-I10/H-2D$^d$/ TAPBPR complex, **(c)** gP18-I10/H-2D$^d$, **(d)** gP18-I10/H-2D$^d$/TAPBPR complex, **(e)** p29/H-2L$^d$/TAPBPR complex, **(f)** QL9/H-2L$^d$/TAPBPR complex, **(g)** QL9/H-2L$^d$/TAPBPR complex.

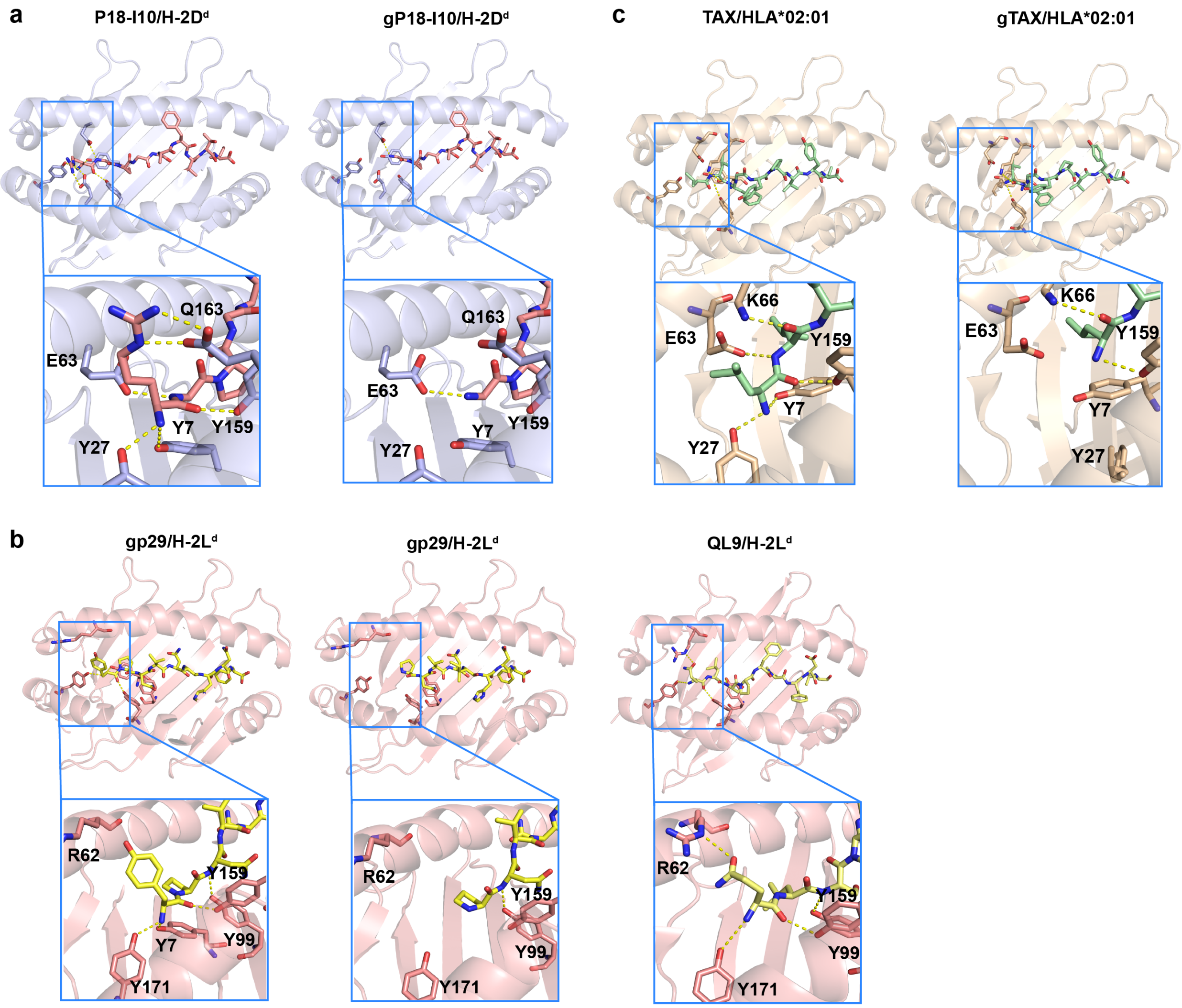


**Supplementary Figure 2. Reduced N-terminal contacts between destabilizing peptides and MHC-I.** Crystal structures of the MHC-I peptide binding groove of **(a)** P18-I10/H-2D$^d$ (PDB 5IVX)[1] and gP18-I10/H-2D$^d$ (modelled from 5IVX), with the bound peptides shown in light red. **(b)** Structure of p29/H-2L$^d$ (PDB 1LD9)[2], gp29/H-2L$^d$ modelled from 1LD9[2] and QL9/H-2L$^d$ (PDB 3TF7)[3], with bound peptides shown in yellow. **(c)** TAX/HLA-A*02:01 (PDB 1DUZ) and gTAX (PDB 1DUY)[4], with the bound peptide shown in green. Insets focus on the N-terminal region of each peptide, with 3 Å polar contacts between the peptide and the indicated MHC-I residues shown as dotted yellow lines.

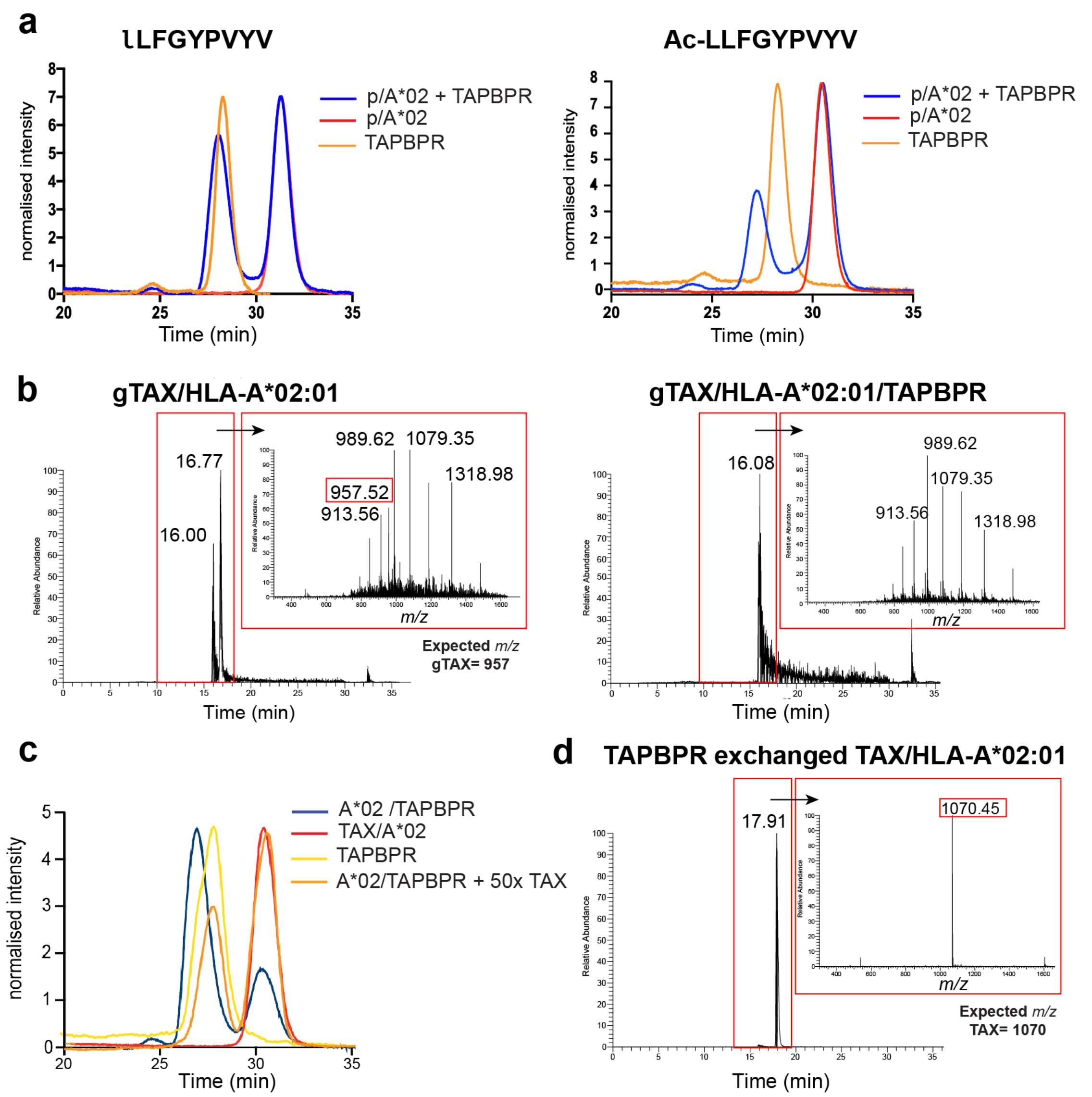


**Supplementary Figure 3. Assessment of peptide occupancy of HLA-A*02:01/TAPBPR complexes. (a)** SEC elution profile of lLFGYPVYV/HLA-A*02:01, where l denotes a D-Leucine residue (left), and Ac-LLFGYPVYV/HLA-A*02:01, where Ac-L denotes an acetylated N-terminal Lecuine residue (right), in the presence or absence of TAPBPR at an equimolar concentration. All binding experiments were performed in the presence of 10 mM GM dipeptide. **(b)** Analysis of peptide occupancy of gTAX/HLA-A*02:01 complex (left) and HLA-

A*02:01/TAPBPR complex (right) by LC-MS. Chromatograms shown are filtered to display ions of interest. Inset: MS analysis of the region indicated. **(c)** SEC analysis of HLA-A*02:01/TAPBPR complex dissociation in the presence of high affinity (TAX) peptide. **(d)** MS analysis of TAX/HLA-A*02:01 isolated from HLA-A*02:01/TAPBPR complexes loaded with TAX. All MS analysis is done on SEC purified HLA-A*02:01.

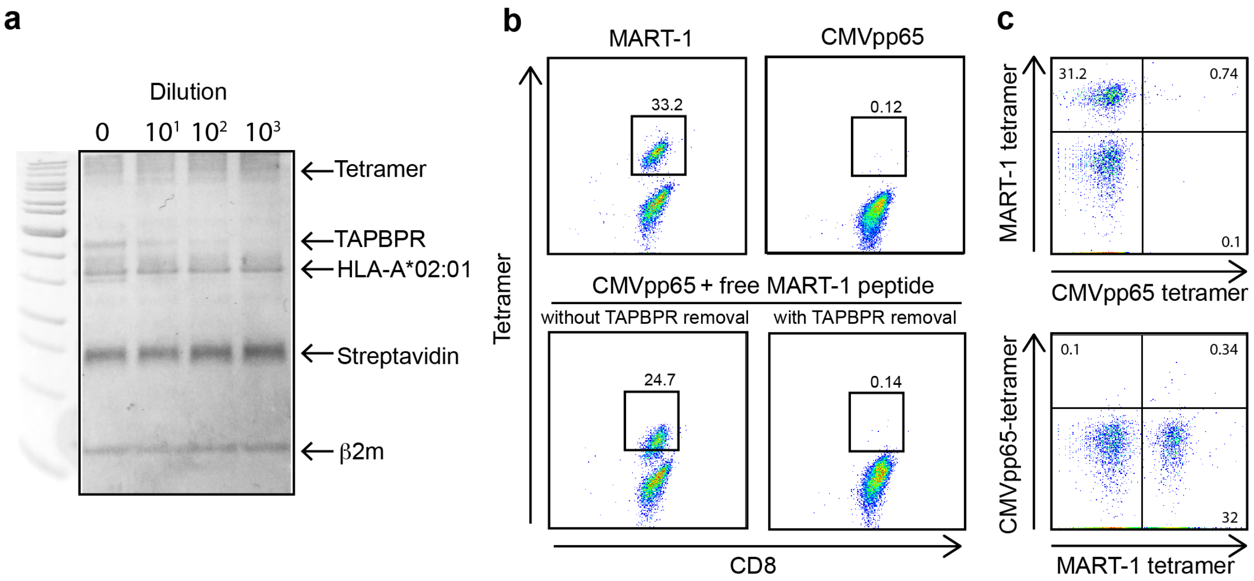


**Supplementary Figure 4. Removal of TAPBPR abrogates the exchange of peptides on and between tetramers for library production. (a)** SDS-PAGE of tetramers washed using a 100 kDa spin filter at 10, 100 and 1,000-fold dilutions. **(b)** Representative plots showing tetramer staining of DMF5 T-cells with HLA-A*02:01 tetramers prepared by TAPBPR exchange of either the MART-1 epitope specific to the DMF5 TCR, or the irrelevant epitope CMVpp65 (top panels). A very low level (0.12%) of CMVpp65-tetramer positive DMF5 T-cells can be detected, likely due to non-specific staining. DMF5 T-cells were independently stained with CMVpp65 tetramers incubated with free MART1 peptide at 10-fold molar excess (relative to pMHC-I) without TAPBPR removal (bottom left) or upon complete removal of TAPBPR, as shown in (a) (bottom right), showing recovery of a low (0.14%) level of background staining in the absence of TAPBPR.

**(c)** Staining of DMF5 T-cells using a 1:1 mixture of PE-CMVpp65:APC-MART-1 tetramers (top panel), or PE-MART-1:APC-CMVpp65 tetramers (bottom panel). In each plot, both tetramer samples were prepared individually using TAPBPR exchange, followed by complete removal of TAPBPR and excess peptide, mixing of the two tetramers and overnight incubation at 4 $^{o}$C. The absence of a significant PE-tetramer positive population (0.1% - x-axis) in the top panel, or an APC-tetramer positive population in the bottom pane (0.1% - y-axis) is indicative of a negligible background of cross-exchange of peptides between tetramers in the absence TAPBPR, allowing their incorporation into stable tetramer libraries. Numbers in the plots indicate the percentage of total cells.

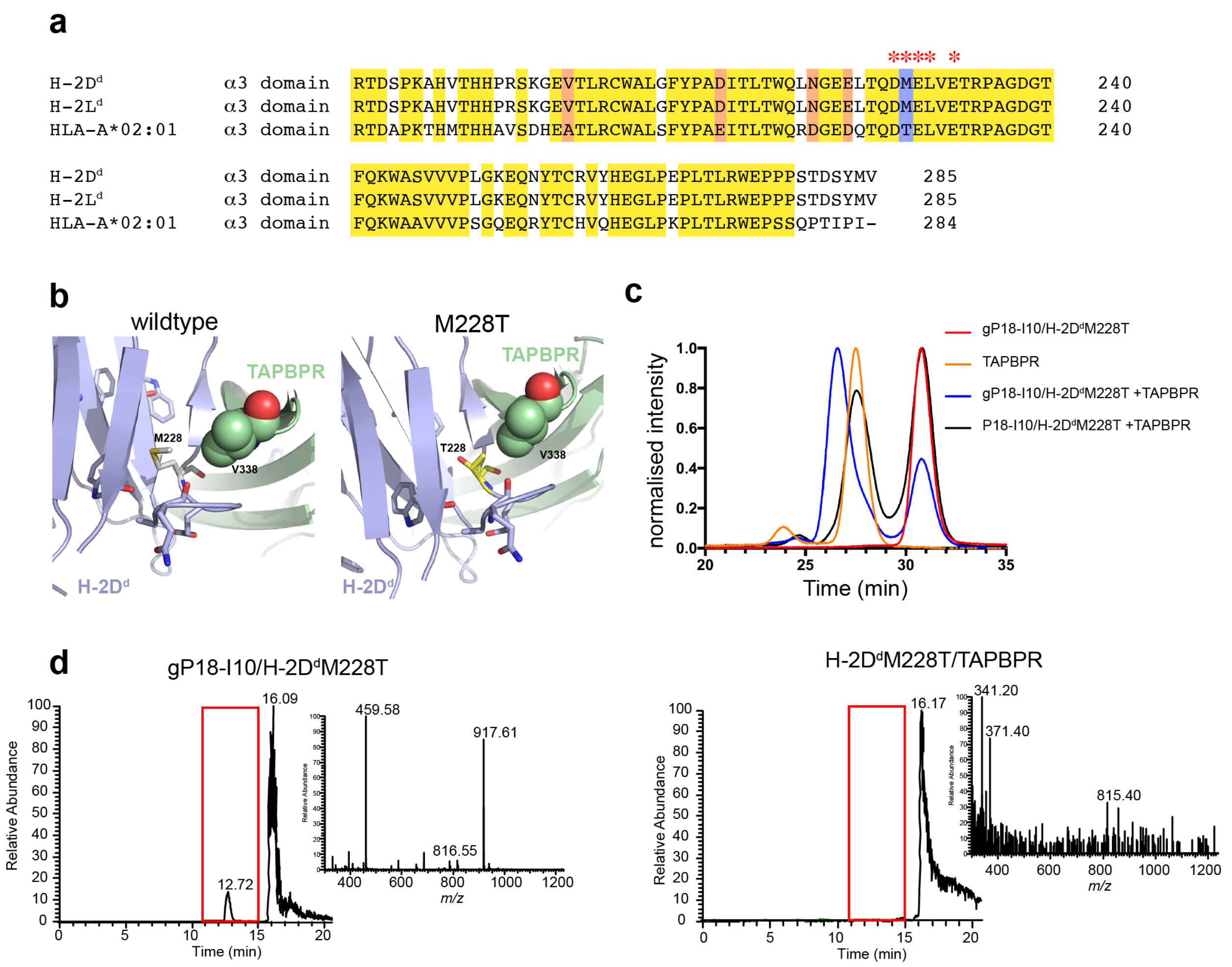


**Supplementary Figure 5. Fine-tuning of MHC-I/TAPBPR interactions towards tetramer library production through the design of α3 domain mutants.** **(a)** Alignment of the α3 domain sequences from murine H-2D$^{d}$, H-2L$^{d}$ and human HLA-A*02:01. Conserved residues are highlighted in yellow, conservative substitutions are highlighted in pale red. The M228T mutation site is highlighted in blue. * indicates residues directly participating in TAPBPR binding, as shown in published mutagenesis studies and crystal structures. **(b)** TAPBPR/H-2D$^{d}$ α3 domain interface from PDB ID 5WER[5] (left panel) and with the M228T mutation modeled (right panel). **(c)** SEC traces of H-2D$^{d}$M228T refolded with either high affinity p18-I10 or goldilocks gP18-I10 peptides, with and without TAPBPR. **(d)** LC/MS peptide occupancy analysis of SEC-purified gP18/H-2D$^{d}$M228T and H-2D$^{d}$M228T/TAPBPR peaks from (c).

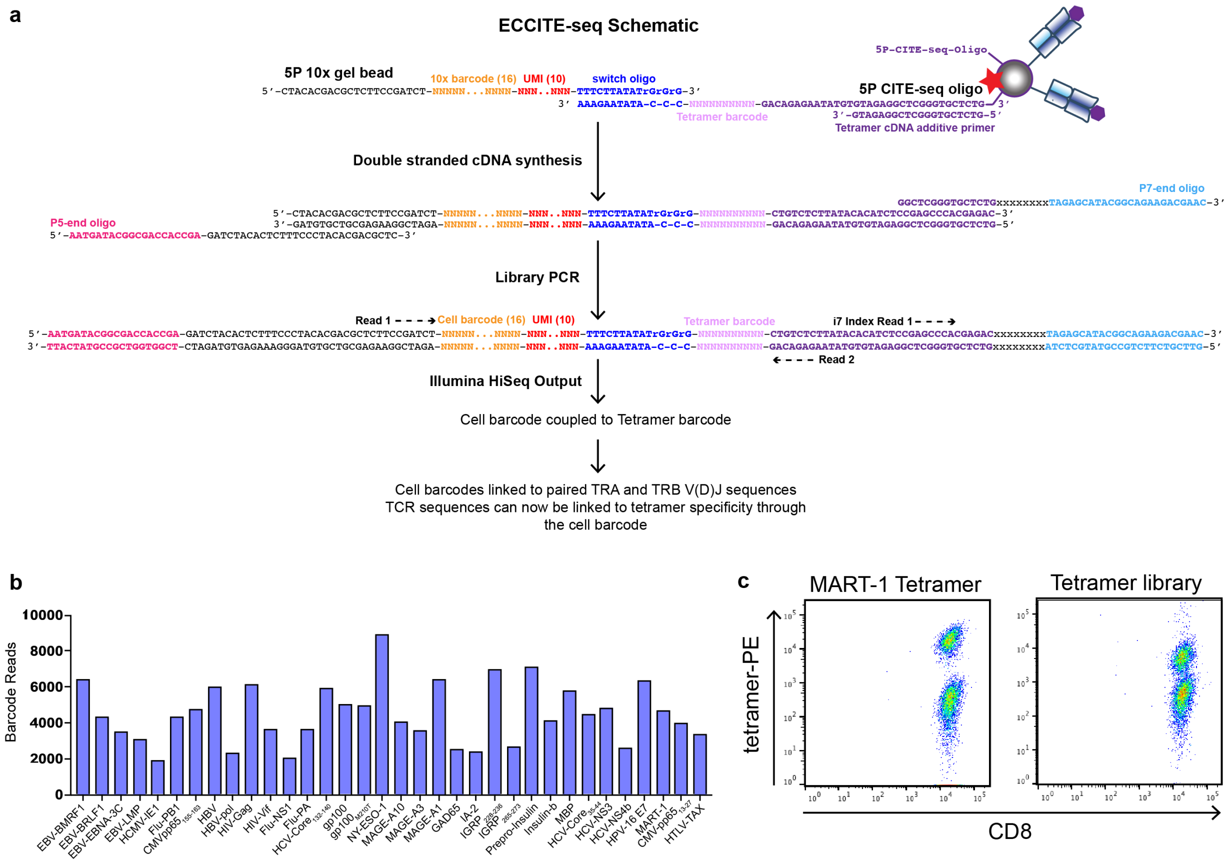


**Supplementary Figure 6: ECCITE-seq adapted to capturing barcoded MHC tetramers.**

**(a)** Biotinylated 5P ECCITE-seq oligos were conjugated to streptavidin tetramers. The oligos contain a unique tetramer barcode, a switch oligo sequence that provides a handle for 10x compatibility and incorporation of 5P 10x gel bead oligos during cDNA synthesis, in addition to a 3’ Illumina NGS sequencing handle (Nextera read 2). The 10x 5P kit was used with specific protocol modifications (as outlined in Online Methods) to capture oligo-derived tags and mRNA-derived cDNA. Only oligo capture is shown here. After separation of the large and small fractions, following cDNA amplification with additive primer, the low molecular weight fraction was amplified with 10x Genomics SI-PCR oligo and a Nextera P7 oligos to create a sequencing library compatible with Illumina instruments. The high molecular weight cDNA fraction was processed according to manufacturer's instructions. Tetramer tags and TCR cDNAs from the same cell will share the same cell barcode and can be associated. **(b)** Bulk amplification of all PE-tetramer barcodes contained in library 2. **(c)** Comparative staining of DMF5 Jurkat T-cells with a single (non-barcoded) PE-tetramer prepared by TAPBPR exchange of the MART-1 peptide (left panel), versus an equal concentration of the full barcoded tetramer library 2 of 34 epitopes, including MART-1 (right panel).

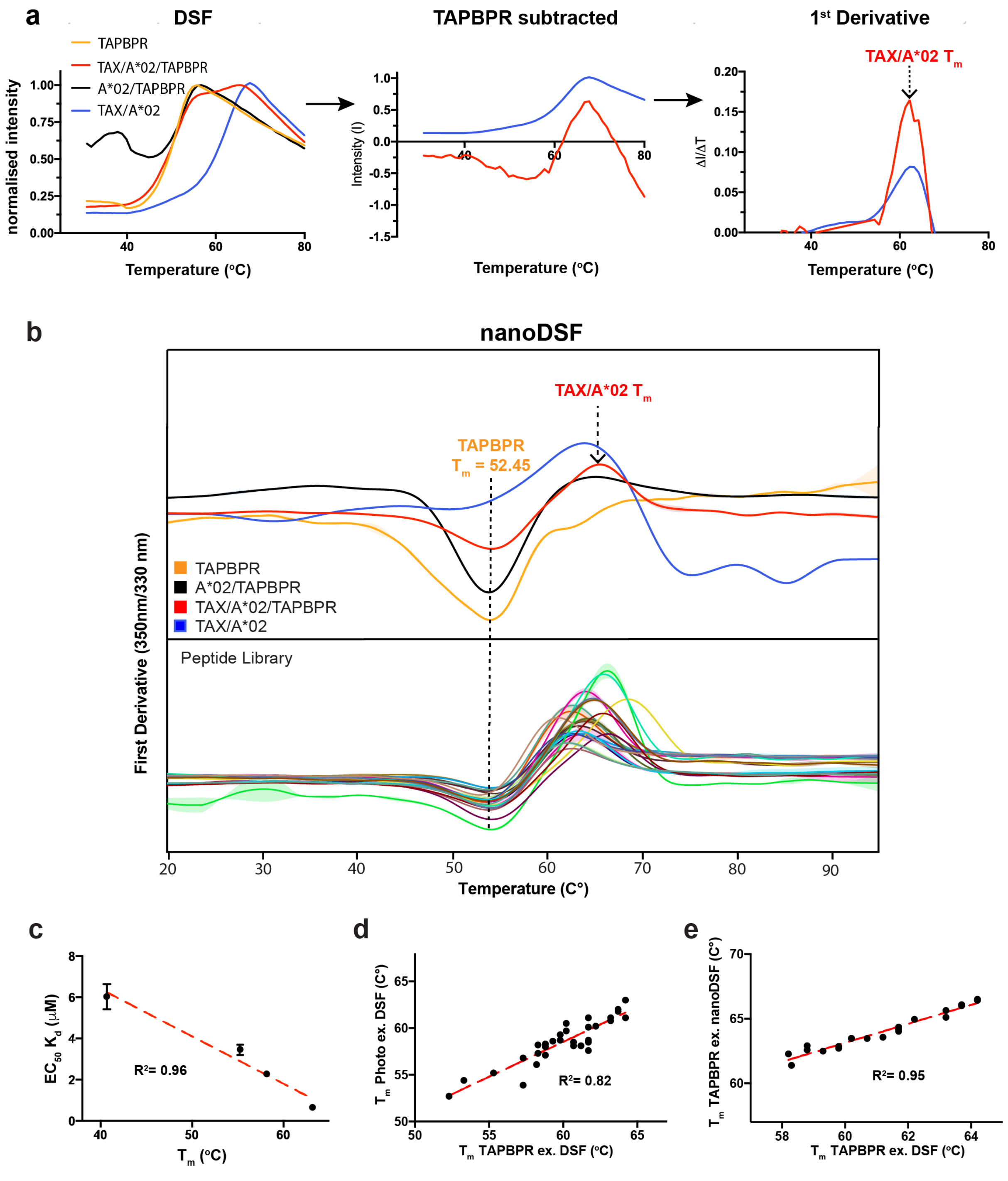


**Supplementary Figure 7. High-throughput validation of peptide loading on HLA-A*02:01/TAPBPR complexes using differential scanning Fluorimetry (DSF). (a)** Derivation of melting temperatures for TAPBPR exchanged TAX/HLA-A*02:01 from conventional DSF data. The DSF trace of TAPBPR alone (left, orange line) was subtracted from that of TAPBPR exchanged HLA-A*02:01 (red line) to obtain the subtracted curve (center, red line). The $T_m$ can be extracted by taking the first derivative (right, red line). As a reference the DSF trace of TAX/HLA-A*02:01 is processed in the manner but without removing the TAPBPR trace (blue line). **(b)** (upper panel) nanoDSF traces showing a negative inflection point for TAPBPR at a $T_m$ of 52.5 °C, and a positive inflection point for TAX/HLA-A*02:01 at a $T_m$ of 66.5 °C. (lower panel) nanoDSF traces of HLA-A*02:01/TAPBPR loaded with different peptides from our library showing a consistent negative inflection point corresponding to the TAPBPR $T_m$, and a positive inflection point corresponding to the $T_m$ values of different pMHC molecules. **(c)** Correlation between $T_m$ values of different pHLA-A*02:01 molecules, measured by DSF, and $EC_{50}$ values of peptide binding on empty HLA-A*02:01/TAPBPR complexes, measured by Bio-Layer Interferometry (Figure 2g). $EC_{50}$ error bars were estimated from 3 independent experiments per peptide. **(d)** Correlation between $T_m$ values of pMHC molecules prepared using photo-exchange and TAPBPR-mediated peptide exchange. **(e)** Correlation between $T_m$ values of TAPBPR exchanged pMHC molecules measured by conventional DSF as shown in (a), and by nanoDSF (b).

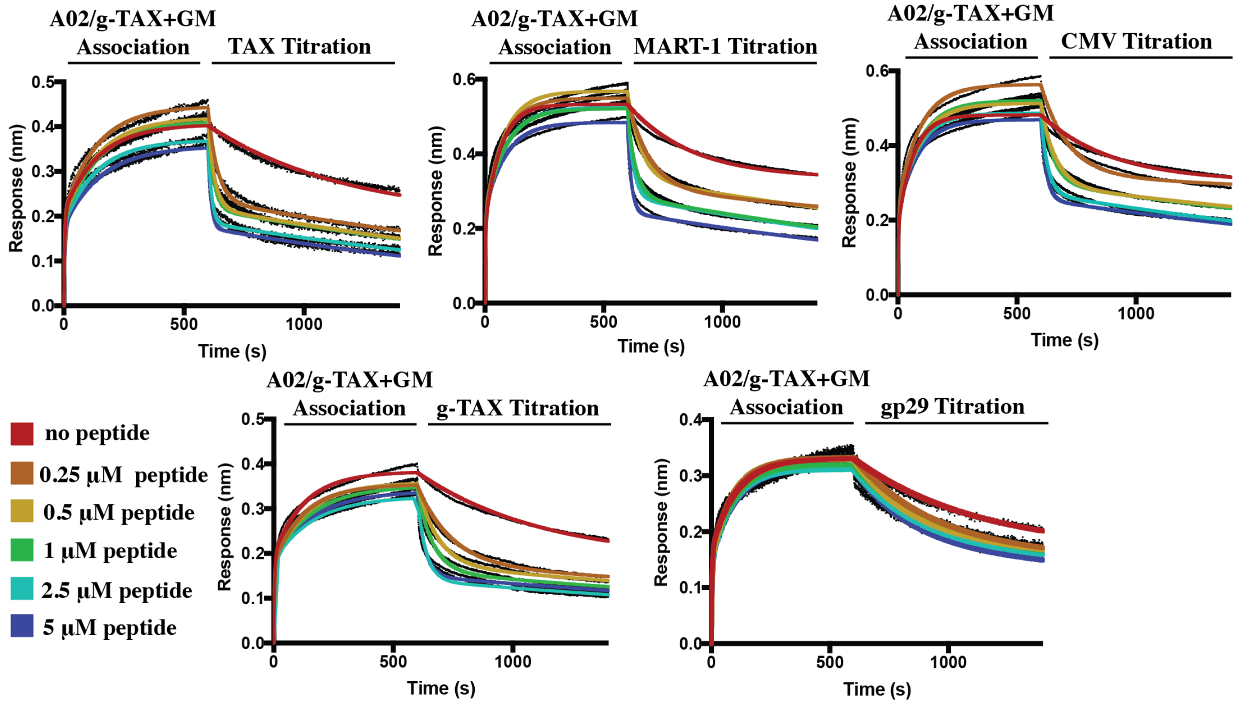


**Supplementary Figure 8: Measurement of peptide-induced HLA-A*02:01 dissociation from TAPBPR using Bio-layer Interferometry.** Binding and dissociation of HLA-A*02:01 from immobilized TAPBPR on a tip surface. The association of a peptide deficient HLA-A*02:01 to TAPBPR was achieved by the addition of 10 mM GM dipeptide (Figure 2c). The dissociation of HLA-A*02:01 was measured in the presence of buffer supplemented with increasing concentrations of the indicated peptide. Dissociation data was corrected using a reference biosensor of immobilized TAPBPR (not bound to HLA-A*02:01), in buffer. Data shown are representative of 2 independent experiments. Raw data points are shown in black. All series were fit locally using a 2:1 model with a $R^2$ of 0.99 or greater, shown as colored curves. The derived pseudo first-order dissociation rate constants, $k_d$, are plotted as a function of peptide concentration in Figure 2g.

| **Peptide index** | **TAPBPR exchange $T_m$ ($^oC$)** | **photo exchange $T_m$ ($^oC$)** | **$IC_{50}$ (nM)** |
|---|---|---|---|
| NB1 | 58.8 ± 0.7 | 57.1 ± 0.1 | 13 |
| NB2 | 61.7 ± 0.7 | 58.7 ± 0.2 | 21 |
| NB3 | 61.7 ± 0.7 | 57.6 ± 0.8 | 9 |
| NB4 | 60.7 ± 0.7 | 58.1 ± 0.1 | 8 |
| NB5 | 59.8 ± 0.7 | 59.3 ± 0.3 | 21 |
| NB6 | 57.3 ± 0 | 53.9 ± 0.3 | 28 |
| NB7 | 60.2 ± 0 | 59.7 ± 0.2 | 30 |
| NB8 | 58.3 ± 0 | 58.2 ± 0.5 | 3 |
| NB9 | 59.8 ± 0.7 | 58.7 ± 0.4 | 4 |
| NB10 | 61.7 ± 0.7 | 58.5 ± 0.8 | 241 |
| NB11 | 63.7 ± 0.7 | 62.0 ± 0 | 6 |
| NB12 | 62.2 ± 1.4 | 60.2 ± 0.3 | 9 |
| NB13 | 58.8 ± 0.7 | 58.0 ± 0.4 | 26 |
| NB14 | 57.3 ± 1.4 | 56.8 ± 0.2 | 96 |
| NB15 | 61.2 ± 0 | 58.1 ± 0.4 | 18 |
| NB16 | 63.2 ± 0 | 61.1 ± 0.3 | 6 |
| NB17 | 64.2 ± 0 | 63.0 ± 0.03 | 3.4 |
| NB18 | 64.2 ± 0 | 61.1 ± 0.5 | 19 |
| NB19 | 61.7 ± 0.7 | 61.1 ± 0.4 | 33 |
| NB20 | 60.2 ± 0 | 60.5 ± 0.1 | 42 |
| NB21 | 58.8 ± 0.7 | 58.3 ± 0.2 | 48 |
| NB22 | 59.3 ± 0 | 58.6 ± 0.1 | 13 |
| NB23 | 60.7 ± 0.7 | 58.5 ± 0.5 | 13 |
| NB24 | 63.2 ± 1.4 | 60.8 ± 0.3 | 3 |
| NB25 | 63.7 ± 0.7 | 61.8± 0.3 | 14 |
| NB26 | 61.7 ± 0.7 | 60.1± 0.1 | 19 |
| NB27 | 53.3 ± 0 | 54.4± 0.2 | 480 |
| NB28 | 52.3 ± 0 | 52.7± 0.2 | 635 |
| NB29 | 58.3 ± 0 | 57.3± 0.1 | 4 |
| MART1 (reference) | 58.2 ± 0 | 56.1± 0.5 | 254 |

**Table 1: pMHC $T_m$ values of Neuroblastoma neoepitopes included in Library 1**. Conventional DSF was performed on pHLA- A*02:01 samples prepared by either TAPBPR-mediated peptide exchange (column 2), or exchange using UV irradiation of a photo-sensitive conditional peptide ligand (column 3), as outlined in Online Methods. A 20-fold molar excess of free peptide was used to promote exchange during a 1 hr incubation at room temperature, for all experiments. DSF profiles were analysed as shown in Supplementary Fig. 7a. All measurements were performed in PBS buffer. Errors represent the standard deviation of 3 replicates, individually analysed. $IC_{50}$ values were obtained from NetMHCpan-4.0 [6].

| Origin | Epitope | Sequence | Tm (°C) | $IC_{50}$ |
|---|---|---|---|---|
| EBV BMRF1 | 208-216 | TLDYKPLSV | 51.3 ± 0 | 36 |
| EBV BRLF1 | 109-117 | YVLDHLIVV | 55.3 ± 0 | 4 |
| EBV EBNA3c | 284-293 | LLDFVRFMGV | 55.3 ± 3.0 | 54 |
| EBV LMP-1 | 159-167 | YLQQNWWTL | 54.3 ± 0 | 9 |
| HCMV IE1 | 81-89 | VLAELVKQI | 50.3 ± 1.0 | 154 |
| Influenza PB1 | 413-421 | NMLSTVLGV | 60.2 ± 0 | 10 |
| HCMV pp65 | 155-163 | QMWQARLTV | 53.3 ± 0 | 81 |
| HBV core | 19-27 | FLFPSDFFPSV | 62.2 ± 0 | 4 |
| HBV Pol | 575-583 | FLLSLGIHL | 53.3 ± 1.0 | 9 |
| HIV Gag | 77-85 | SLYNTVATL | 50.8 ± 0.5 | 54 |
| HIV Vif | 101-109 | GLADQLIHL | 60.2 ± 0 | 10 |
| Influenza NS1 | 1122-1130 | AIMDKNIIL | 52.8 ± 0.5 | 54 |
| Influenza PA | 70-78 | ALLKHRFEI | 53.3 ± 0 | 37 |
| HCV Core | 132-140 | DLMGYIPAV | 52.3 ± 0 | 10 |
| gp100 | 209-217 | IMDQVPFSV | 54.3 ± 0 | 6 |
| gp100 | 209-217 (210T) | ITDQVPFSV | 50.3 ± 0 | 102 |
| NY-ESO-1 | 157-165 | SLLMWITQA | 51.3 ± 0 | 26 |
| MAGE-A10 | 254-262 | GLYDGMEHL | 53.3 ± 0 | 9 |
| MAGE-A3 | 271-279 | FLWGPRALV | 53.3 ± 0 | 12 |
| MAGE-A1 | 278-286 | KVLEYVIKV | 61.7 ± 0.5 | 7 |
| GAD65 | 114-122 | VMNILLQYV | 47.3 ± 0 | 23 |
| IA-2 | 805-813 | VIVMLTPLV | 47.8 ± 0.5 | 72 |
| IGRP | 228-236 | LNIDLLWSV | 53.8 ± 0.5 | 243 |
| IGRP | 265-273 | VLFGLGFAI | 50.3 ± 0 | 16 |
| Prepro-insulin | 15-24 | ALWGPDPAAA | 53.3 ± 0 | 234 |
| Insulin b | 10-18 | HLVEALYLV | 61.2 ± 0 | 7 |
| MBP | 110-118 | SLSRFSWGA | 47.8 ± 0.5 | 24 |
| HCV core | 35-44 | YLLPRRGPRL | 53.3 ± 0 | 245 |
| HCV NS3 | 1406-1415 | KLSGLGINAV | 60.2 ± 0 | 41 |
| HCV NS4b | 1807-1816 | LLFNILGGWV | 55.3 ± 0 | 49 |
| HPV16 E7 | 12-20 | MLDLQPETT | 43.9 ± 0.5 | 1812 |
| CMV pp65 | 496-503 | NLVPMVATV | 55.3 ± 0 | 25 |
| HTLV | 11-19 | LLFGYPVYV | 61.2 ± 0 | 3 |
| MART1 (reference) | 26-35 | ELAGIGILTV | 58.2 ± 0 | 254 |

**Table 2: pMHC $T_m$ values of viral, tumor, autoimmune epitopes included in Library 2**. Conventional DSF was performed on HLA-A*02:01/TAPBPR complexes following 1 hr incubation with each listed peptide. A 20-fold molar excess of free peptide was used to promote exchange during a 1 hr incubation at room temperature, for all experiments. DSF profiles were analysed as shown in Supplementary Fig. 7a. Measurements were performed in PBS buffer supplemented with 0.5% DMSO to increase peptide solubility. Errors represent the standard deviation of 3 replicates, individually analysed. $IC_{50}$ values were obtained from NetMHCpan-4.0 [6].

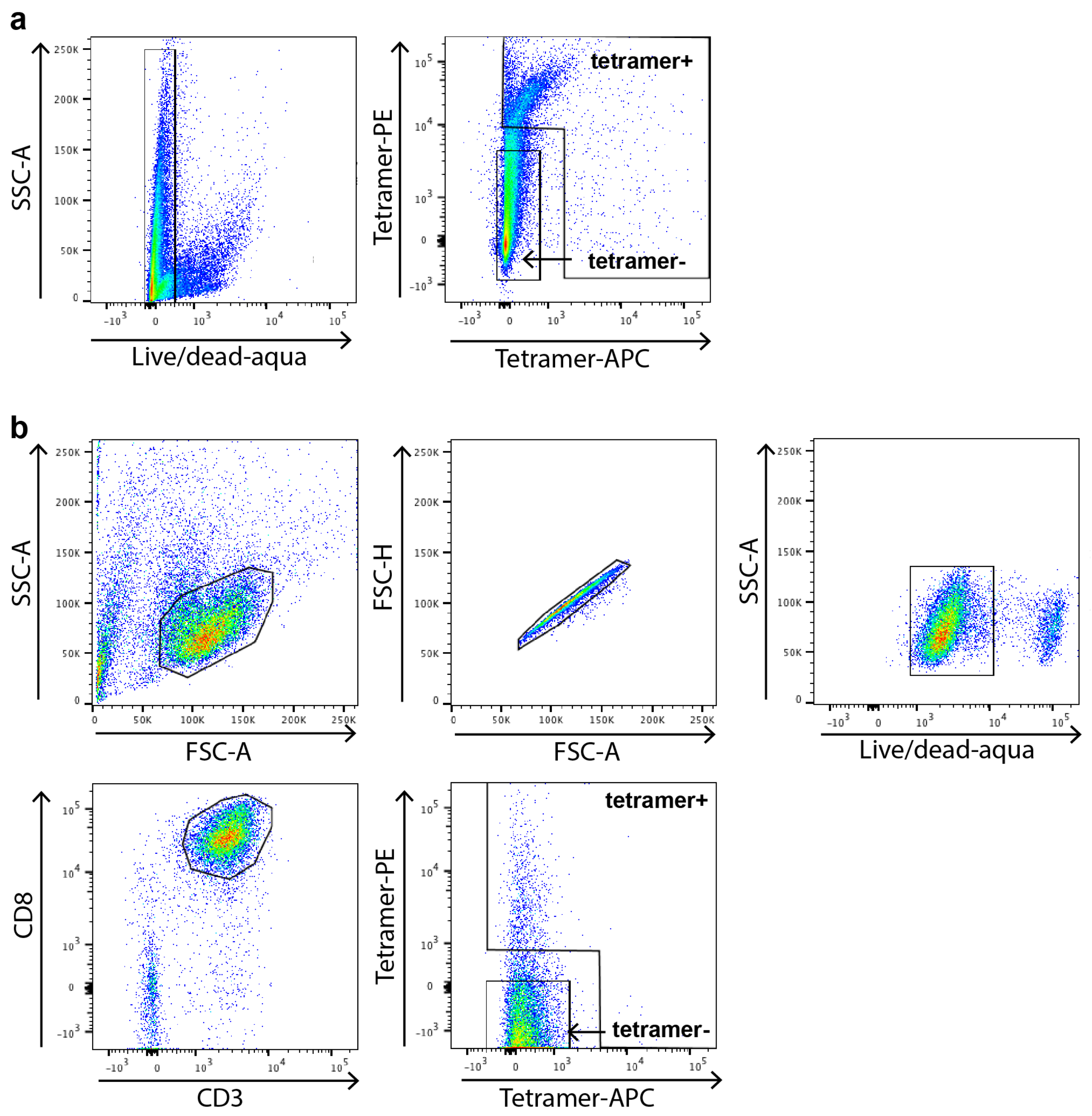


**Supplementary Figure 9. Flow cytometry gating strategy. (a)** Gating strategy used to sort tetramer positive cells expanded *in vitro* with NB epitope pulsed DCs. Mixed cell cultures of pulsed DCs and PBMCs were stained using both PE/APC versions of tetramer library 1, and sorted on a flow cytometer by gating on live cells, then on tetramer positive cells. **(b)** Gating strategy used to sort CD8-enriched splenocytes stained with PE/APC versions of library 2. Cells were sorted by SSC-A and FSC-A to select for lymphocytes, then filtered for single cells using FSC-H and then live cells with LIVE/DEAD-Aqua. From this, CD8+ T-cells were gated based on CD8 and CD3 staining and then tetramer positive cells were collected for sequencing experiments. Due to low observed staining on the APC fluorophore channel for both samples shown in (a,b) >99% of collected cells correspond to the PE-tetramer positive fractions.
